# How to validate a Bayesian evolutionary model

**DOI:** 10.1101/2024.02.11.579856

**Authors:** Fábio K. Mendes, Remco Bouckaert, Luiz M. Carvalho, Alexei J. Drummond

## Abstract

Biology has become a highly mathematical discipline in which probabilistic models play a central role. As a result, research in the biological sciences is now dependent on computational tools capable of carrying out complex analyses. These tools must be validated before they can be used, but what is understood as validation varies widely among methodological contributions. This may be a consequence of the still embryonic stage of the literature on statistical software validation for computational biology. Our manuscript aims to advance this literature. Here, we describe and illustrate good practices for assessing the correctness of a model implementation, with an emphasis on Bayesian methods. We also introduce a suite of functionalities for automating validation protocols. It is our hope that the guidelines presented here help sharpen the focus of discussions on (as well as elevate) expected standards of statistical software for biology.

## Introduction

The last two decades have seen the biological sciences undergo a major revolution. Critical technological innovations such as the advent of massive parallel sequencing and the accompanying improvements in computational power and storage have flooded biology with unprecedented amounts of data ripe for analysis. Not only has intraspecific data from multiple individuals allowed progress in fields like medicine and epidemiology (e.g., The 1000 Genomes Project Consortium, 2015; Human Microbiome Project Consortium, 2012; Neafsey et al., 2015), population genetics (e.g., Lynch, 2007; Lack et al., 2016; de Manuel et al., 2016) and disease ecology (e.g., Rosenblum et al., 2013; Bates et al., 2018), but now a large number of species across the tree of life have had their genomes sequenced, furthering our understanding of species relationships and diversification (e.g., Pease et al., 2016; Kawahara et al., 2019; Upham et al., 2019). Almost on par with with data accumulation is the rate at which new computational tools are being proposed, as evidenced by journals entirely dedicated to method advances, methodological sections in biological journals, and computational biology degrees being offered by institutions around the world.

One extreme case is the discipline of evolutionary biology, on which we focus our attention. While it could be said that many decade-old questions and hypotheses in evolutionary biology have aged well and stood up the test of time (e.g., the Red Queen hypothesis, Van Valen 1973; Lively 1987; Morran et al. 2011; Gibson and Fuentes-González 2015; the Bateson-Dobzhansky-Muller model, Dobzhansky 1936; Muller 1940; Hopkins and Rausher 2012; Roda et al. 2017), data analysis practices have changed drastically in recent years, to the point they would likely seem exotic and obscure to an evolutionary biologist active forty years ago. In particular, evolutionary biology has become highly statistical, with the development and utilization of probabilistic models now being commonplace.

Models are employed in the sciences for many reasons, and fall within a biological abstraction continuum (Servedio et al., 2014), going from fully verbal, highly abstract models (e.g., Van Valen 1973), through proof-of-concept models that formalize verbal models (e.g., Maynard Smith 1978; Reinhold et al. 1999), to models that interact directly with data through explicit mathematical functions (Yule, 1924; Felsenstein, 1973; Hasegawa et al., 1985; Hudson, 1990). Within the latter category, probabilistic models have seen a sharp surge in popularity within evolutionary biology, in conjunction with computational tools implementing them.

Despite the increasing pervasiveness of probabilistic models in the biological sciences, tools implementing such models show large variation not only with respect to code quality (from a software engineering perspective), but also to the provided evidence for correctness (Darriba et al., 2018). This is unsurprising given the challenges in funding software research (Siepel, 2019), and the multidisciplinary nature of method development. Much of the relevant information regarding good coding and statistical practices is out of reach of the average computational biologist, as it is spread across a variety of specialized sources, often obfuscated by its technical and theoretical presentation. The bioinformatics community is thus in dire need of synthetic and accessible resources that provide guidance for code improvement and validation.

Here, we summarize best practices in probabilistic model validation for method developers, with an emphasis on Bayesian methods. We execute two different validation protocols on variations of a simple phylogenetic model, discuss the results, and expand on how to interpret other potential outcomes. We further introduce a suite of methods for automating these protocols within the BEAST 2 platform (Bouckaert et al., 2019). Lastly, we propose method development guidelines for new model contributions, for researchers and reviewers who expect new software to meet not only a desirable standard, but also a reasonable one.

## Probabilistic models

Probabilistic models mathematically formalize natural phenomena having an element of randomness. This is done through probability distributions describing both the observed empirical data – seen as the result of one or more random instantiations of the modeled process – as well the model parameters, which abstract relevant, but usually unknown aspects of the phenomenon at hand. In the domain of evolutionary biology specifically, the historical, stochastic, and highly dimensional nature of evolutionary processes makes the utility of probabilistic models self-evident.

The central component of a probabilistic model, Pr(*D* = *d*|Θ = *θ*), allows us to describe the probability distribution over the data (*D*) given the model parameters (Θ). This probability mass function (pmf; or its continuous counterpart, the probability density function, pdf, *f*_*D*_(*d*|Θ = *θ*)) is sometimes referred to as the likelihood function. Just for this section, we will abuse and simplify notation, and drop variable subscripts, e.g., we will write *f*_*D*_(*d*|Θ = *θ*) as *f* (*d*|*θ*). As illustrated in the next sections, probabilistic models can be hierarchical, in which case there may be several likelihood functions. In a frequentist statistical framework, *f* (*d*|*θ*) is the sole component of an inferential procedure, and is maximized across parameter space during parameter estimation and model comparison.

In the present study we focus on Bayesian inference, where a probabilistic model *ℳ* defines a posterior probability distribution for its parameters,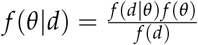. Here, our prior inferences or beliefs about the natural world – represented by the prior distribution *f* (*θ*) – are confronted with and updated by the data through the likelihood function. *f* (*d*) = _Θ_ *f* (*d*|*θ*) *f* (*θ*)*dθ*, the probability of the data, is also known as the marginal likelihood or the model evidence. Crucially, a Bayesian model includes a prior, *f* (*θ*): when models are compared, for example, *f* (*θ*) needs to be taken into account when computing the model evidence *f* (*d*).

Models routinely used in evolutionary biology are often characterized by continuous parameters, and are normally complex enough to preclude analytical solutions for the posterior density *f* (*θ*|*d*), mainly due to the intractability of the integral appearing in the denominator – i.e., the marginal likelihood. In those cases, one can make use of the fact that *f* (*d*) is a constant with respect to the parameters that can be ignored (i.e., *f* (*θ*|*d*) ∝ *f* (*θ*|*d*) *f* (*θ*)), and use techniques like Markov chain Monte Carlo (MCMC) to sample the posterior distribution. This is because MCMC is usually implemented in the form of the Metropolis-Hastings (Metropolis et al., 1953; Hastings, 1970) algorithm, which only requires the posterior to be evaluated up to a constant.

In practice, the Metropolis-Hastings algorithm samples the posterior distribution (also referred to as the “target” distribution) by means of a transition mechanism. If the proposal distribution generated by this mechanism is irreducible, positive recurrent, and aperiodic, and the resulting chain is long enough, then the sampled posterior distribution will closely approximate the target distribution *f* (*θ*|*d*) (Smith and Roberts, 1993; Tierney, 1994; Gelman et al., 2013).

We will spend time considering MCMC in particular, as it is the commonly chosen technique for obtaining samples from *f* (*θ*|*d*) under an implementation of model *ℳ*. A thorough validation effort thus entails verifying the correctness of (i) the model (i.e., *f* (*d*|*θ*) *f* (*θ*)), and (ii) the components involved in the MCMC transition mechanism. We note that the latter are not part of the model, however, and it is possible to sample *f* (*θ*|*d*) with other techniques such as importance sampling, Laplace approximations (Rue et al., 2009), or even by converting the sampling problem into an optimization one (e.g., Zhang and Matsen, 2019).

Finally, we stress that we are interested in practices for verifying model *correctness*. There are other tests and diagnostics employed to ensure that a particular MCMC analysis is converging as expected. Ascertaining whether one or more independent Markov chains have converged to a given posterior distribution is not a correctness test, as that distribution might be very different from the target distribution. We refer the reader interested in these and related topics to Warren et al. (2017), Fabreti and Höhna (2022), Magee et al. (2023) and references therein.

## Validating a Bayesian model

In this section we discuss procedures for validating an implementation of a Bayesian model ℳ. Whenever necessary, we will differentiate between a model implemented as a simulator (S[ℳ]) and as a tool for inference (I[ℳ]). Both S[ℳ] and I[ℳ] must be inspected in order to validate a model ℳ.

### Validating the simulator, S[ℳ]

When a new probabilistic model ℳ is introduced an its inferential engine, I[*ℳ*] – what users employ in empirical analyses – is implemented for the first time, validating I[*ℳ*] requires that a simulator for *ℳ*, S[*ℳ*], be devised and itself validated. A simulator conventionally requires a parameter value as input (i.e., a value for Θ, *θ*, where *θ* might represent the values of more than one parameter), or a prior distribution on those values, *f*_Θ_(*·*). Note that we use “*·*” when referring specifically to the generative function, rather than the value it takes given input. The simulator then outputs a sample of random variable(s), which for hierarchical models will include not only an instantiation *d* of data D, but also the parameters in Θ.

In the case of hierarchical models, it is sometimes useful to consider S[*ℳ*] as a collection of component simulators, each characterized by a different sampling distribution. For example, S[*ℳ*] for model we will work with below (Fig. 1; Table 1) can be seen as an ensemble comprised by:

1. S[*f*_Θ_(*·*)] (where Θ = {*T, Λ,R, **γ***_**0**_},which jointly simulates θ = { *τ, λ,r,**y***_**0**_}
2. S[ *f*_Φ|*T*,Λ_(*·*|*T* = *τ*, Λ= *λ*)], which simulates a Yule tree *ϕ* given an origin age value *τ* and a *λ* (the birth-rate) simulated in (1),
3. S[ *f*_*Y*|Φ,*R*,***Y*0**_ (*·*|Φ = *ϕ, R* = *r*, ***Y***_**0**_ = ***y***_**0**_)], which simulates an array with *n* continuous-trait values (one value per species), ***y***, given a phylogeny *ϕ* with *n* species, an evolutionary rate *r* and ancestral character values ***y***_**0**_ (simulated in (1) and (2), respectively).

Being able to isolate the building blocks of a hierarchical model simulator helps divide and conquer the validation task, especially when some but not all of the sampling distributions are well-known parametric distributions, or when they result from well characterized stochastic processes (see below).

**Figure 1:**
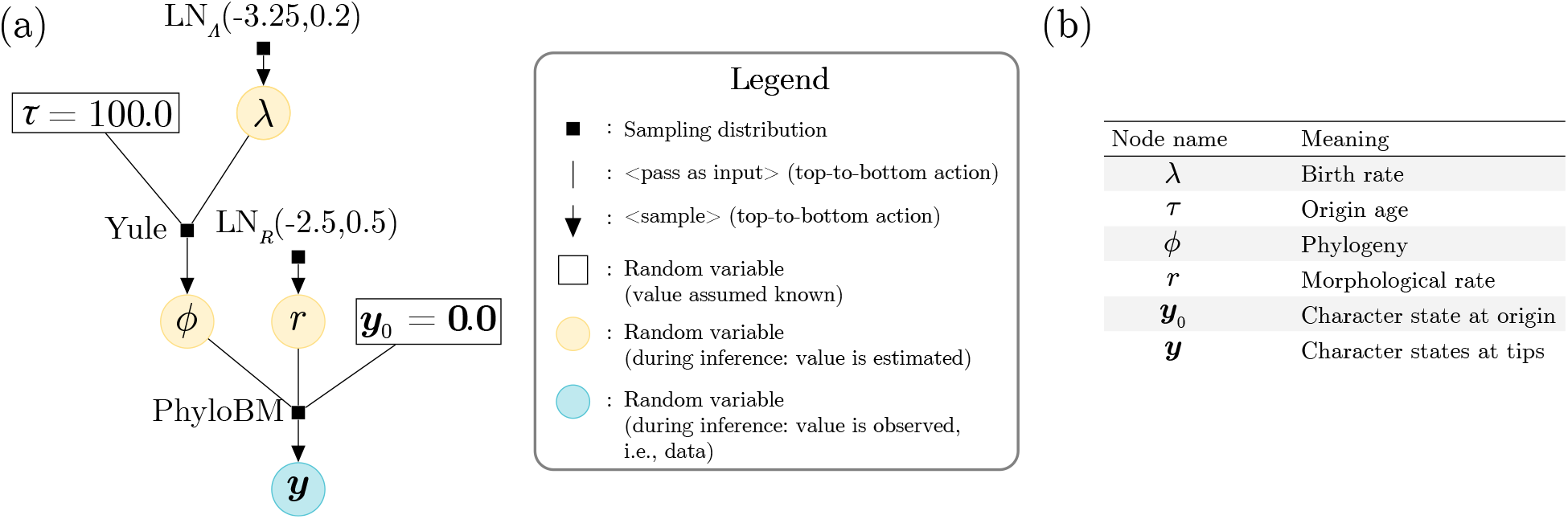
A simple probabilistic graphical (Bayesian) model we will validate in this work. (a) When read from top to bottom, the graphical model describes a generative process (see the legend for the meaning of vertical lines and downward-pointing arrows). If read from bottom to top, the graphical model describes the process of inference (assuming arrows having opposite orientation denoting the flow of information); in this case, the blue and yellow circles represent the data and the parameters being estimated, respectively. A random variable within a rectangular box signifies a parameter whose value is assumed known by the user; these are normally nuisance hyperparameters, or parameters that are not of immediate interest perhaps because they have been estimated elsewhere. (b) Each random variable node in the model, and how they should be interpreted. Table 1 presents more detail on each of the sampling distributions. Briefly, “LN” stands for log-normal, “Yule” for a Yule process also known as a pure-birth model, and “PhyloBM” stands for a phylogenetic Brownian motion model.

**Table 1:**
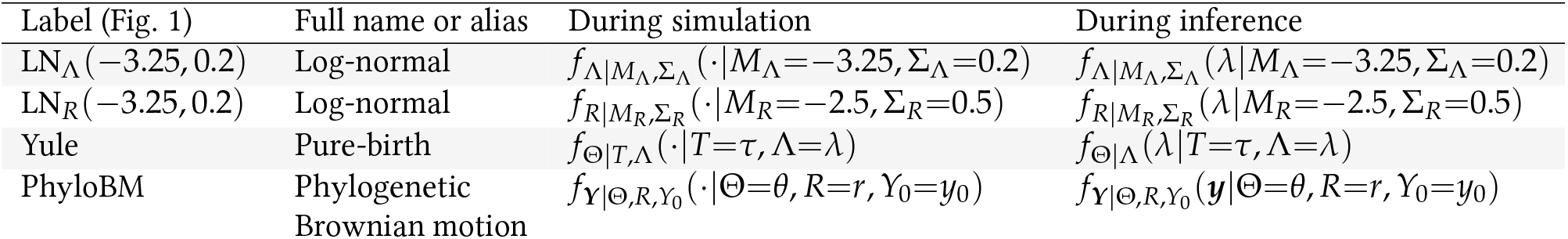
Sampling distributions used in the probabilistic model validated in this work (Fig. 1). Columns “During simulation” and “During inference” specify how the sampling distributions should be read and interpreted, following the notation in the main text.

One way to validate a probabilistic model simulator is by using it to produce (sample) a large number of data sets given a set of parameters. These parameters can be seen as characterizing an ensemble of the entities being modeled. For each data set, one can then construct *α ×* 100%-confidence intervals (where *α ∈* (0, 1) gives the confidence level) for certain summary statistics (e.g., mean, variance, covariance). If the simulator is behaving as expected, one should be able to verify that the (ensemble’s or “true”) summary statistic is contained approximately *α*% of the time within their *α ×* 100%-confidence intervals. An example is the Yule model (also known as the pure-birth model; Yule 1924), a continuous-time Markov process that has been classically employed in phylogenetics to model the number of species in a clade (Yule, 1924; Aldous, 2001). Under a Yule process with a species birth rate of *λ*, the expected tree height, E[*t*_root_], for a tree with *n* tips is:

#### Box 1

**Models characterized by well-known parametric distributions**

One commonly used model in macroevolution for the study of continuous traits is the phylogenetic Brownian motion model (“PhyloBM” in Fig. 1; Felsenstein 1973). The pdf characterizing this model’s sampling distribution is in fact the pdf of the multivariate normal (MVN) probability distribution:

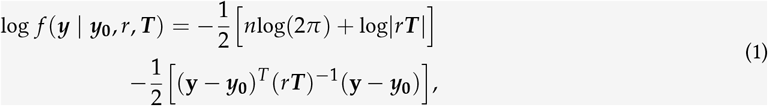

where ***y*** corresponds to the observed values of a trait scored for *n* species, ***y***_**0**_ is the trait value at the root of the tree, *r* is the instantaneous rate of change (i.e., the evolutionary rate, and sometimes represented by *σ*^2^), and *r****T*** is the variance-covariance matrix. ***T*** is a matrix whose elements are deterministically defined by tree *ϕ*’s topology and branch lengths; see Fig. 1 below).

The probability density function in equation (1) describes the distribution that would result from an infinite number of BM “experiments” (each experiment being non-mean-reverting, and representing an independent evolutionary trajectory). Under this model, *θ* = *{****y***_**0**_, *r*, ***T*** *}* and *d* = *{****y****}* (but note that sometimes researchers treat *ϕ* and consequently ***T*** as data).

##### Validating a phylogenetic BM simulator

The MVN is a well-characterized parametric distribution. When used as the sampling distribution of the phylogenetic BM process, it explicitly defines the expected trait value for each species (***y***_**0**_), as well as their trait value variances and covariances. The latter comes from the variance-covariance matrix; for the tree shown in Figure 2 and with *r* = 0.1, this matrix is:

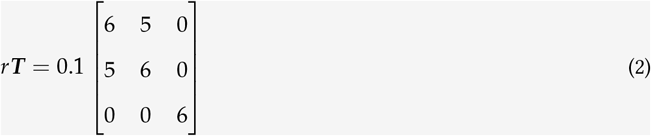

Together, variance-covariance matrix *r****T*** and ***y***_**0**_ = [0.0, 0.0, 0.0] characterize a population of phylogenetically related species trait values whose means are 0.0, variances are 6.0, and co-variances are 5.0 (between species “A” and “B”) and 0.0 (between species “C”and either “A” or “B”).

**Figure 2:**
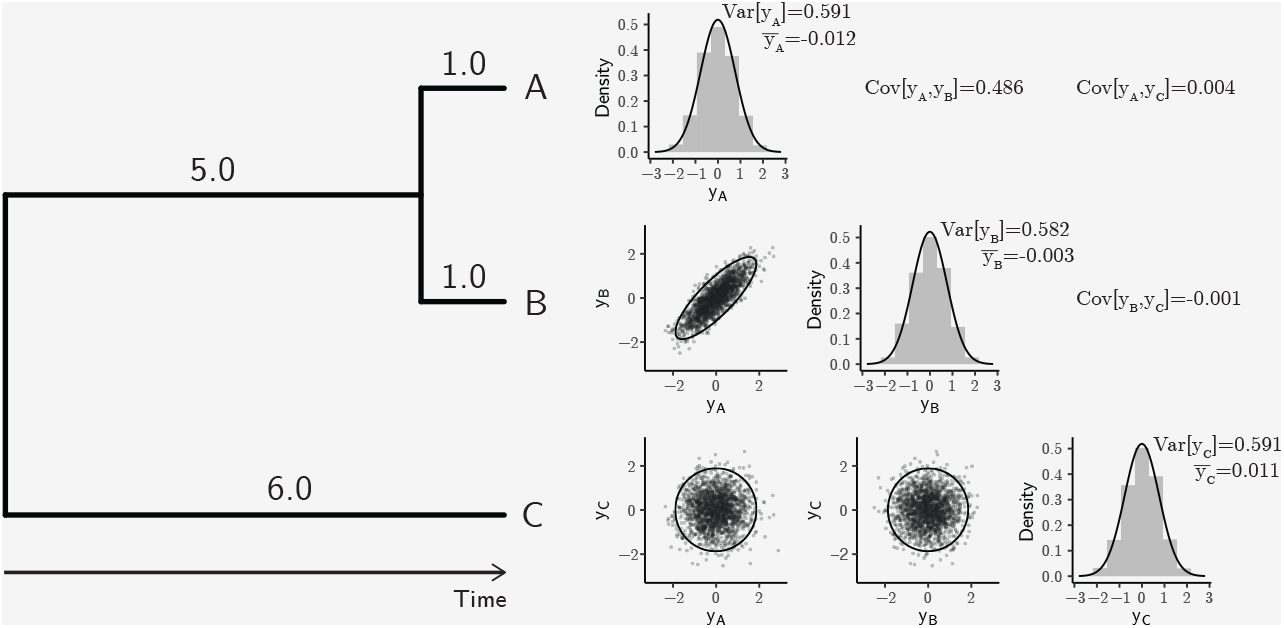
A sample of 1000 draws from a MVN distribution, each representing the evolutionary trajectory of one continuous trait along the species tree on the left. The root trait value, ***y***_**0**_, and the evolutionary rate of the process, *r*, were set to 0.0 and 0.1, respectively. The panel on the right shows histograms of 1000 trait values sampled from the MVN for each species, as well as their covariation.

Figure 1 shows the distributions of trait values and their variances and covariances for one sample of one thousand independent realizations of phylogenetic BM processes. One can see that the sample’s average trait value and the variances and covariances approach their expectations. More rigorously, one can follow the method described in the main text and verify that those expectations fall within their 95%-confidence intervals 95% of the time, as calculated from a large number of samples (Supplementary Fig. 1 and Supplementary Table 1).

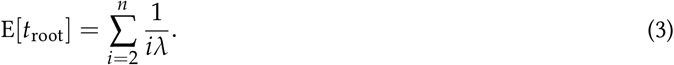

One can then verify, for example, if E[*t*_root_] is 95% of the time within *±*1.96 standard errors of the average Yule-tree height (from each sampled data set). Confirming that this is the case indicates S[ *f*_Φ|Λ_(*·*|*T* = *τ*, Λ= *λ*)] is correctly implemented (Fig. 3). In Box 1, we illustrate this procedure for the (parametric) sampling distribution underlying the phylogenetic Brownian motion model (“PhyloBM”; Felsenstein, 1973). Protocols for validating I[ℳ] (see below) will also normally validate S[ℳ] at the same time.

**Figure 3:**
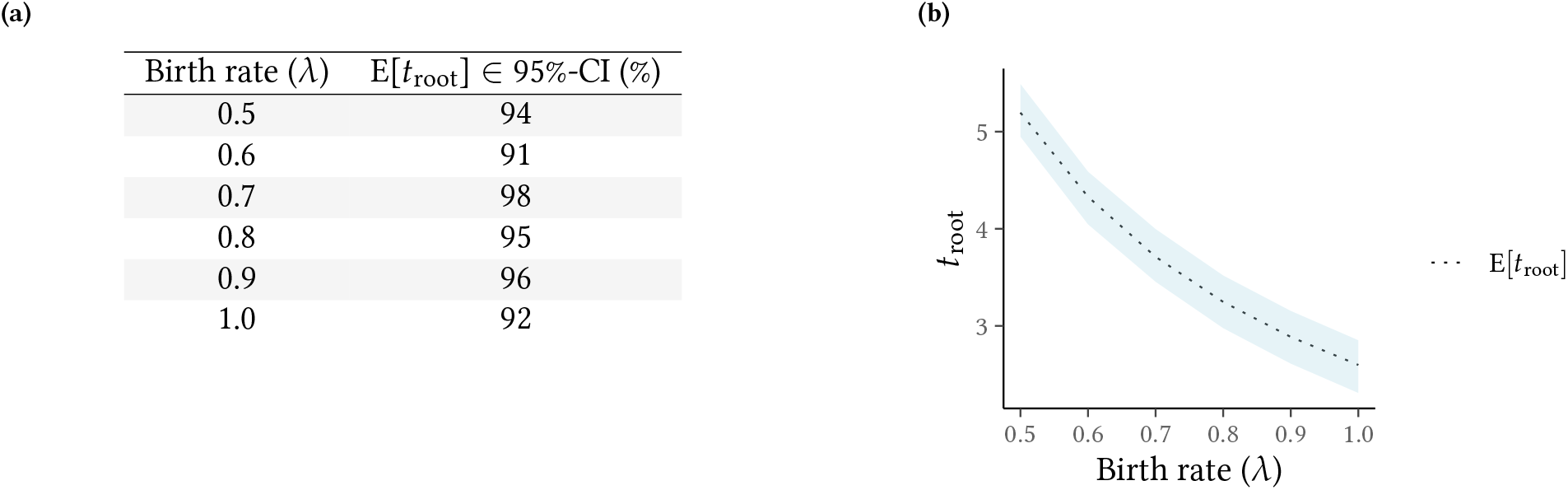
Validation of Yule tree simulator. (a) Number of simulated data sets (out of 100) for which the expected tree height (*t*_root_) was inside the 95%-confidence interval about its sample average – if the simulator is correct, we expect this number to be between 90 and 99 about 95% of the time. Each data set consisted of 50 twenty-taxon simulated Yule trees. (b) The area shaded in light blue represents the 95%-confidence interval about the average tree height, obtained from the 5,000 Yule trees simulated in (a). Simulations were carried out with the TreeSim R package (Stadler, 2011).

We note that we have so far used S[ℳ] to represent a *direct* simulator under model ℳ (Table 2), meaning each and every sample generated by S[ℳ] is independent. This is contrast with other simulation strategies, such as conducting MCMC under model ℳ with no data (i.e., “sampling from the prior”), given specific parameter values (*θ*). This latter approach may be the only option if S[ℳ] has not been yet implemented, and it is predicated upon the existence of correct implementations of both an inferential engine I[*ℳ*]^*’*^ and of proposal functions. We distinguish I[ℳ]^*’*^ from I[ℳ] because simulations are being carried out precisely to validate I[ℳ]. Unless MCMC simulations are done with I[ℳ]^*’*^ – an independent implementation of I[ℳ] – they can introduce circularity to the validation task.

**Table 2:**
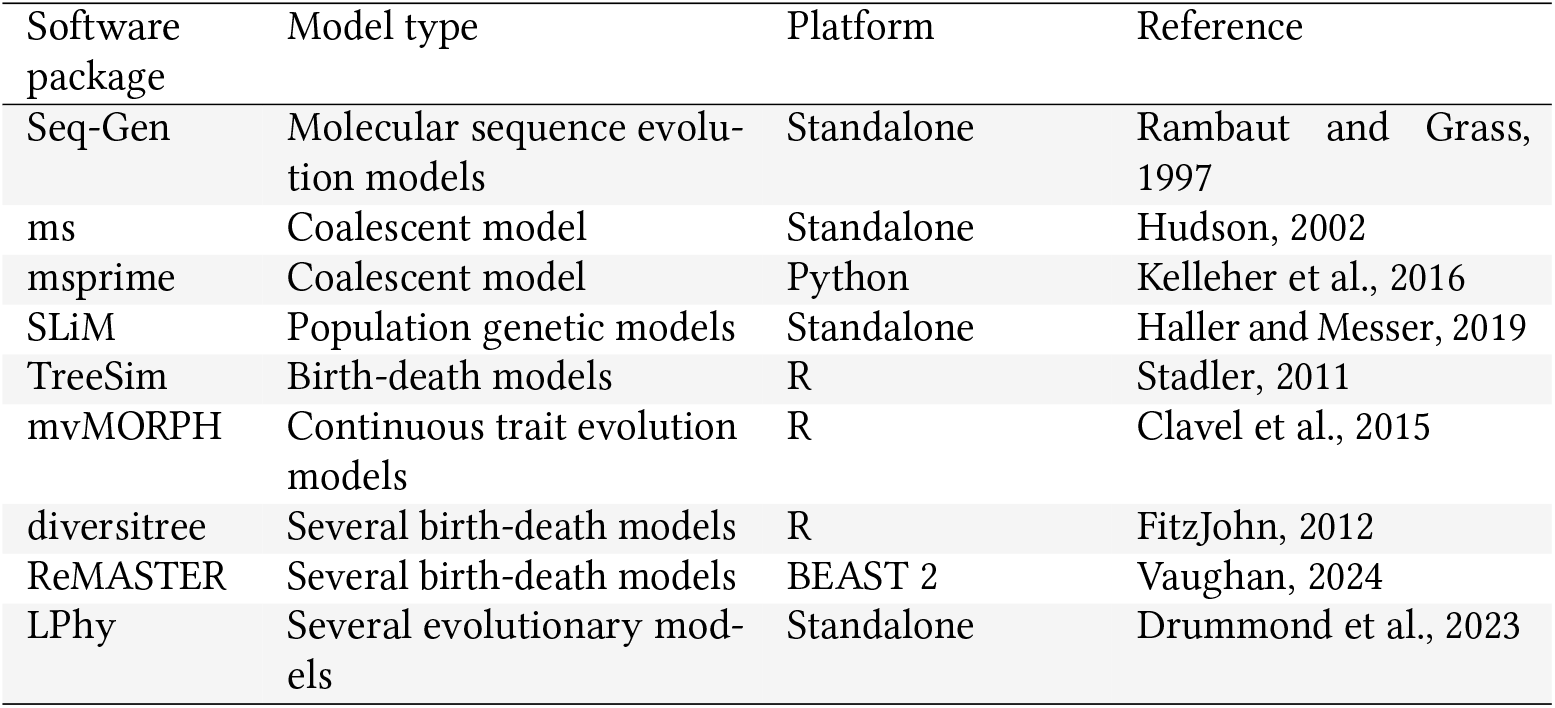
A non-exhaustive list of direct simulation software used for various models in evolutionary biology.

| Software package | Model type | Platform | Reference |
| --- | --- | --- | --- |
| Seq-Gen | Molecular sequence evolution models | Standalone | Rambaut and Grass, 1997 |
| ms | Coalescent model | Standalone | Hudson, 2002 |
| msprime | Coalescent model | Python | Kelleher et al., 2016 |
| SLiM | Population genetic models | Standalone | Haller and Messer, 2019 |
| TreeSim | Birth-death models | R | Stadler, 2011 |
| mvMORPH | Continuous trait evolution models | R | Clavel et al., 2015 |
| diversitree | Several birth-death models | R | FitzJohn, 2012 |
| ReMASTER | Several birth-death models | BEAST 2 | Vaughan, 2024 |
| LPhy | Several evolutionary models | Standalone | Drummond et al., 2023 |

## Validating the inferential engine, I[ℳ]

The more complex the natural phenomenon under study, the more difficult it will be to strike a good balance between model practicality and realism (Levins, 1966). The popular aphorism rings true: “all models are wrong but some are useful” (Box, 1979). Very simple models are easier to implement in efficient inference tools, but will commonly make assumptions that are likely to be broken by the data. Conversely, complex models will fit the data better, but may become unwieldy with increasing levels of realism.

A large number of parameters can cause overfitting and weak identifiability, and inference under highly complex models might be prohibitively slow (Shapiro et al., 2000). Deciding on the utility of a model for real-world problems is a daunting task (Brown and Thomson, 2018; Shepherd and Klaere, 2018), and is a challenge we do not address in the present contribution. Such model appraisals (what we call “model characterization” below) are normally carried out after a model is published, often in multiple contribution bouts, and are critical for a model’s longevity. Analyses of model fit against data are normally accompanied by discussions on assumption validity, and more rarely by benchmarking and scrutinization of model behavior and implementation (e.g., Maddison et al., 2007; Stadler, 2010; Rabosky et al., 2013; Rabosky and Goldberg, 2015; Moore et al., 2016).

When a new model *ℳ* is initially proposed, however, authors must ensure that their methods can at the very least robustly recover generating parameters. In this section, we discuss a few techniques that can be employed to assess the correctness of a parameter-estimation routine. These techniques assume that one can accurately simulate from a probabilistic data-generating process (see previous section).

### Coverage validation

Our discussion on how to ensure a Bayesian model is well-calibrated and thus correct will mostly follow the ideas in Cook et al. (2006) and Talts et al. (2018). The basic idea is presented in the flowchart in figure 4 (acquamarine dotted box and what is above it), and consists of three stages: simulation, inference, and coverage calculation. Once we have a validated simulator for model ℳ, we start by sampling *n* parameter sets ***θ*** = *{θ*_*i*_ : 1 *≤ i ≤ n}* from its prior, *f*_Θ_(*·*), i.e.:

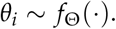

For each parameter set *θ*_*i*_, we then sample a data set *d*_*i*_ from *f*_*D*|Θ_(*·*|Θ = *θ*_*i*_):

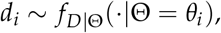

These two steps conclude the “simulation” stage of this validation protocol. With ***d*** = *{d*_*i*_ : 1 *≤ i ≤ n}*, we use the inferential machinery I[ℳ] under evaluation to compute *f*_Θ|*D*_(*θ*_*i*_|*D* = *d*_*i*_) for each *d*_*i*_. Recall that we assume the posterior distribution defined by *f*_Θ|*D*_(*θ*|*D* = *d*) over Θ will be approximated with MCMC, an algorithm that generates a large sample of size *L* of parameter values from that posterior distribution, 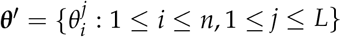. At this point, we have concluded the inference stage of this validation pipeline.

**Figure 4:**
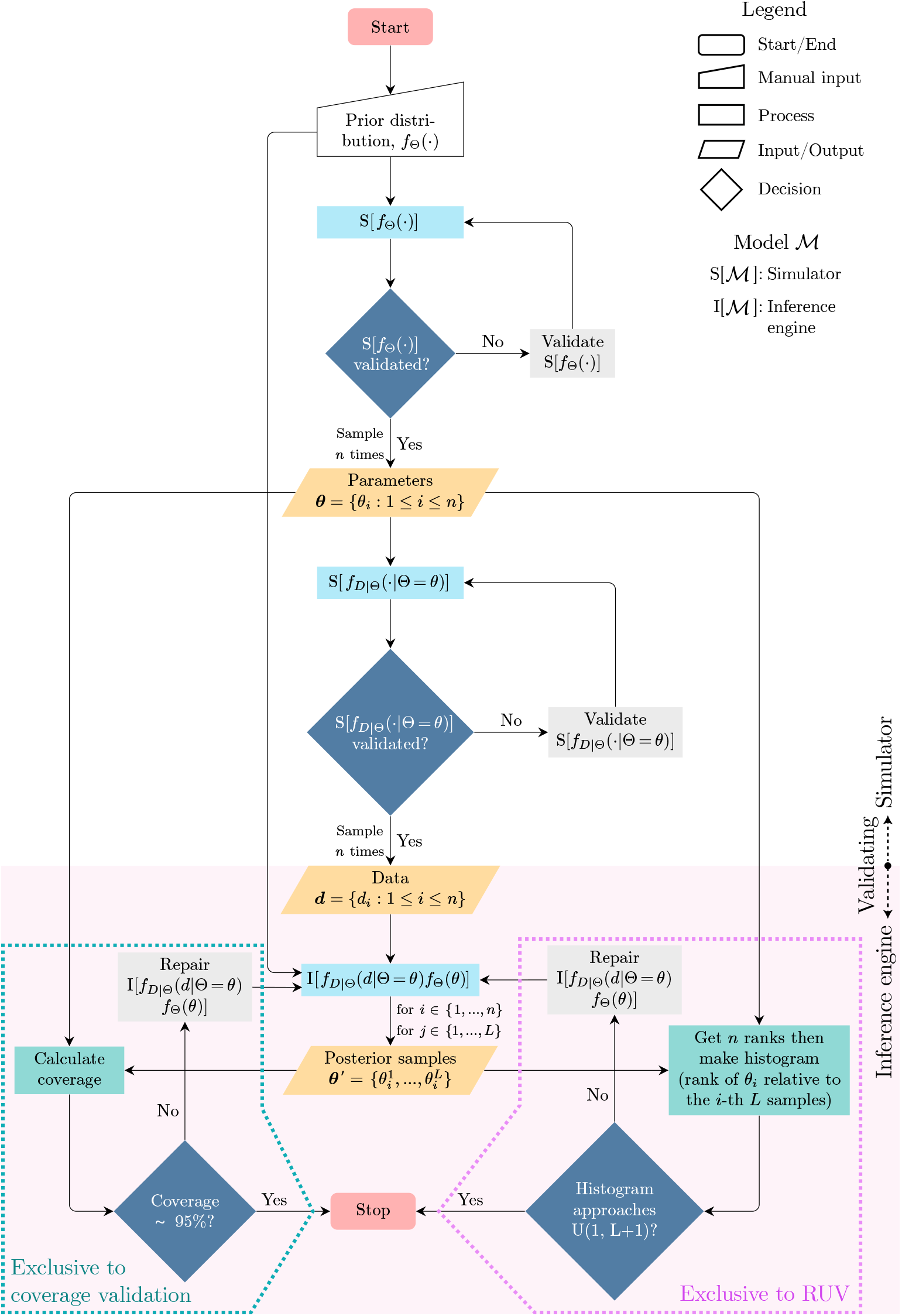
Flowchart of the validation of a Bayesian model. Standard flowchart symbols are explained in the legend. The flowchart area with a clear background is where (true) parameters and data are generated, and where the model simulator(s) is validated. The flowchart area shaded in pink mark the steps involved in validating the inference engine once the data has been generated. ***θ*** denotes a vector with *n* elements, where each element is an i.i.d. parameter(s) sample from its (their) corresponding prior(s) *f*_Θ_(). Analogously, ***d*** denotes a vector with *n* elements, where each element is an i.i.d. data sample from the corresponding likelihood(s) *f*_*D*| Θ_( | Θ = *θ*). ***θ***, holds *n × L* elements, with each being one of the *L* posterior samples for each of the *n* parameter samples in ***θ***. All *L* posterior samples obtained from the *i*-th data set *d*_*i*_ comprise together what one would call the posterior distribution over *θ*_*i*_. Posterior samples are commonly obtained through MCMC. U(*l, u*) denotes a uniform distribution with and including lower and upper bounds *l* and *u*, respectively. The acquamarine dotted box encloses the stages of the pipeline that are exclusive to the coverage validation procedure. The pink dotted box encloses the stages of the pipeline that are exclusive to the rank-uniformity validation (RUV) procedure.

The third stage and final stage consists of investigating coverage properties of uncertainty intervals. The critical expectation here is that if the inferential engine is correct, we will be able to obtain interval estimates with precise coverage properties. More concretely, let us first define the highest posterior density (HPD) interval. For a credibility level *α ∈* (0, 1), we define *I*_*α*_(*d*) := (*a*(*d, α*), *b*(*d, α*)) such that:

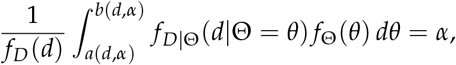

where *f*_*D*_(*d*) is a constant that can be ignored. By defining Cred(*I*_*α*_(*d*)) = *α*,

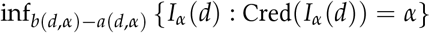

yields the shortest interval with the required credibility. Note that we approximate a particular *I*_*α*_(*d*_*i*_) from the *i*-th *L* samples obtained with MCMC, in ***θ***.

Now taking a set of parameter values *θ*_*i*_ sampled from *f*_Θ_(*·*) it can be shown that Pr (*θ*_*i*_ *∈ I*_*α*_(*d*)) = *α*, i.e., that 100 *× α*% HPDs have nominal coverage under the true generative model (a proof is provided in the supplementary material). More formally, the coverage of *n* intervals obtained as above will be distributed as binomial random variable with *n* trials and success probability *α*. When *n* = 100 and *α* = 0.95, the 95%-central interquantile interval for the number of simulations containing the correct data-generating parameter is between 90 and 99 (Table 3). If we ascertain that I[ℳ] of a Bayesian model produces coverage lying within the expected bounds, we say the model has passed the coverage validation, and is well-calibrated and correct.

**Table 3:** The 95% central interquantile intervals for the number of HPD intervals covering the true parameter value (obtained during coverage validation), under different credibility levels and numbers of replicates. Assuming model correctness, the number of true simulated values that fall within their corresponding 100 *× α*%-HPDs (coverage) is binomially distributed with *n* trials and probability of success *α*.

| Credibility level % ($100 \times \alpha$) | $n$ (replicates) | Lower quantile | Upper quantile |
| --- | --- | --- | --- |
| 90 | 100 | 84 | 95 |
|  | 200 | 171 | 188 |
|  | 500 | 436 | 463 |
| 95 | 100 | 90 | 99 |
|  | 200 | 184 | 196 |
|  | 500 | 465 | 484 |
| 99 | 100 | 97 | 100 |
|  | 200 | 195 | 200 |
|  | 500 | 490 | 499 |

At this point, we will take a moment to remark that the usefulness of model coverage analysis in Bayesian inference is only manifest when *θ*_*i*_ is sampled from *f*_Θ_(*·*). Method developers may be tempted, for example, to calculate coverage for specific parameter values – perhaps chosen across a grid over parameter space – using a different prior during inference. In such cases, we emphasize that obtaining a coverage lower than 95% (for 95% HPDs) does not necessarily mean that a model is incorrectly implemented; conversely, obtaining exactly 95% coverage does not imply model correctness. Coverage values only have bearing on model correctness if, and only if, random variables are sampled from the same prior distribution used in inference.

We provide examples of coverage validation attempts in Figure 5, which shows coverage graphical summaries for data simulated under the model represented in Figure 1. This model is deliberately simple for the sake of brevity and clarity in the discussion below. The parameters in this model are the phylogenetic tree Φ, the species birth rate Λ, and the continuous-trait evolutionary rate *R* (we assume the continuous trait value at the root, ***Y***_**0**_, is known and set it to **0.0** for all simulated data sets). When the model is correctly specified between simulation and inference (“Scenario 1”, Fig. 5a), coverage is close to 95% and adequate for both Λand *R*, which indicates that I[ℳ] – as implemented in BEAST 2, the software we used – is well-calibrated and correct.

**Figure 5:**
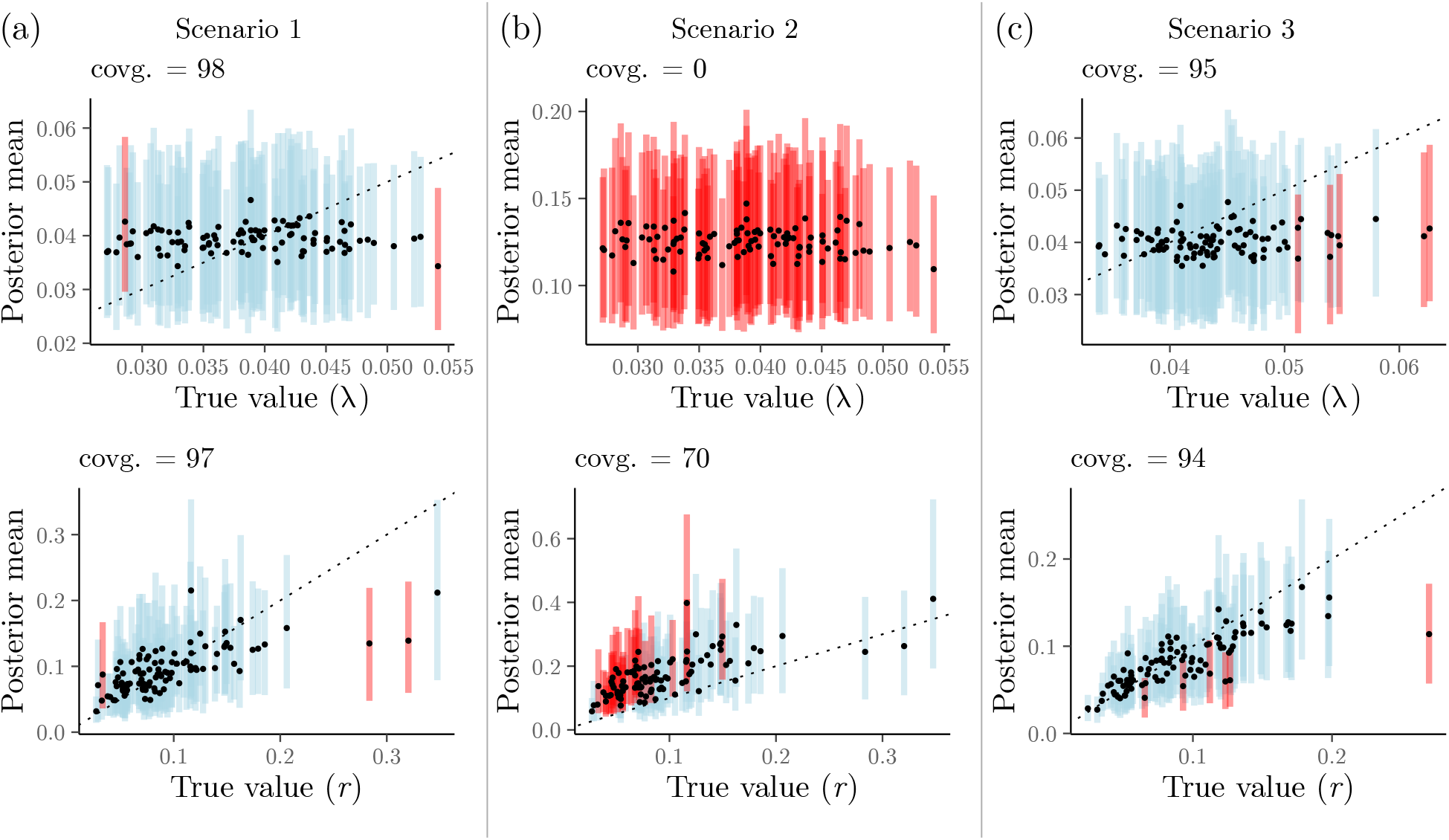
Coverage validation analyses of the Bayesian hierarchical model in Fig. 1. Panels show the true (i.e., simulated) parameter values plotted against their mean posteriors (the dashed line shows *x* = *y*). Dots and lines (100 per panel) represent true values and their 95%-HPDs, respectively. Simulations for which 95%-HPDs contained the true value are highlighted in blue, otherwise are presented in red. (a) In “Scenario 1”, the model was correctly specified, and we simulated trees with 3 to 300 taxa using rejection sampling (approximately one in ten trees were rejected). (b) In “Scenario 2”, the model was incorrectly specified in inference (see main text), and we used the same data sets simulated in “Scenario 1”. (c) In “Scenario 3” the model was specified just as in “Scenario 1”, with the difference that rejection sampling was substantially greater (we rejected a large number of trees, approximately 90%, keeping those having between 100 to 200 tips).

In “Scenario 2” of Figure 5 (Fig. 5b), however, we misspecify the model during inference, setting the prior distribution on Λto be a log-normal with a mean of -2.0 (rather than -3.25, as specified in the simulation procedure; Fig. 1). In contrast with scenario 1, coverage is 0.0 for Λand 70% for *R*, both much lower and outside the expected coverage bounds (Table 3). These numbers indicate that one or more of the parts comprising model ℳused in I[ℳ] differs from their counterparts in S[ℳ]. This result was expected because we purposefully made the models in simulation and inference differ; we know I[ℳ] is correct because of the results from scenario 1. Of course, in a real-world validation experiment the model should be correctly specified, and such a result would suggest a problem with the inferential machinery (provided the the simulator had been previously validated).

Lastly, in “Scenario 3” of Figure 5 (Fig. 5c), we specified the model just like in scenario 1, but carried out substantial rejection sampling during simulation. Approximately 90% of all simulated trees were rejected based on their taxon count; trees were rejected if they had fewer than 100 or more than 200 taxa. As with scenario 1, coverage fell within the expected ranges for a correct model implementation. This result may strike the reader as odd: if I[ℳ] expects trees with a wide range of tip numbers, and we feed it simulated trees within a narrow tip number interval, should this not lower coverage? For example, one may have expected the estimated *λ* to be consistently higher or lower than the true *λ*. Λis nonetheless challenging to infer under the current model, as suggested by estimates falling around the corresponding prior mean value; unlike in scenario 2, however, here the prior mean parameter was correctly specified. As a result, coverage validation was not capable of detecting any symptoms arising from the rejection of tree samples.

Scenario 3 brings home the point that an incorrect model implementation may pass coverage validation, unless model misspecification is sufficiently severe (e.g., scenario 2), or parameter estimate location is highly responsive to the evidence in the data – unlike Λin the examined model. (In the supplement we expand on this point using a different and simpler model, and show that, if extreme, rejection schemes will be detected by coverage validation as a model misspecification issue; Supplementary Table 1.) Put simply, obtaining appropriate coverage may not be enough to ascertain that a model is correct. Potential biases in parameter estimates may remain undetected unless more investigation is done (see rank-uniformity validation below).

The three scenarios we explored above illustrate how coverage validation results can be interpreted in terms of model implementation correctness. One can additionally capitalize on this validation setup and gauge how accurate our inferential tool can be for different parameters. The easier it is to estimate a parameter, the higher should be the correlation between its posterior mean and its generating “true” value. In our scenarios 1 and 3, the species birth rate Λwas hard to estimate given the sizes of the phylogenetic trees. Conversely, the continuous-trait evolutionary rate, *R*, was more easily identifiable, as revealed by the higher correlation between its true values and their posterior means. We conclude this section by noting that the absence of correlation between parameter estimates and their true values (sometimes referred to as “weak unidentifiability”) should not be taken as a sign that a model is incorrect – inappropriate coverage values should.

### Rank-uniformity validation (RUV)

Talts et al. (2018) showed that one can devise other tests that might be more powerful to detect problems than just looking at the coverage of Bayesian HPD intervals. In particular, given ***θ*** = *{θ*_*i*_ : 1 *≤ i ≤ n}* (produced according to the protocol in Fig. 4), those authors demonstrated (Theorem 1 therein) that if the inference machinery I[ℳ] works as intended, the distribution of the rank *r*_*i*_ of *θ*_*i*_ relative to 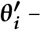 – i.e., the rank of the *i*-th parameter value relative to its corresponding *L* MCMC chain samples – will follow a uniform distribution on [1, *L* + 1] (Fig. 4, pink dotted box; Fig. 6a). In other words, if one were to sort all true parameter values *θ*_*i*_ against 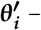 – their corresponding *L* MCMC posterior samples – the first (smallest ranking) 10% out of *n θ*_*i*_ values should account for approximately 10% of the total rank mass; the next 10% of (higher ranking) *θ*_*i*_ values should account, again, for approximately 10% of the total rank mass, and so on.

**Figure 6:**
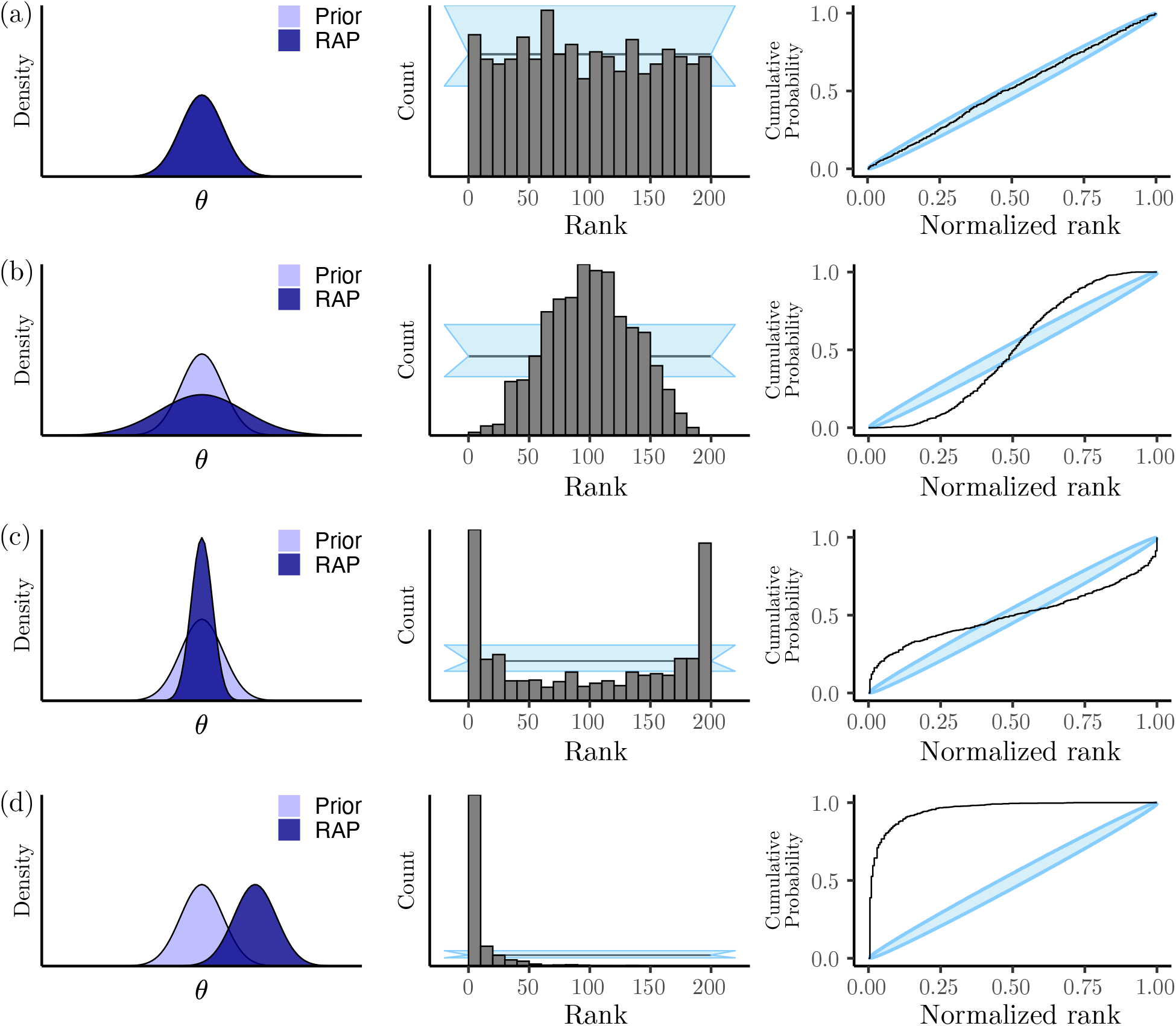
Patterns observable after inference in rank-uniformity validation (RUV). We explain how to interpret the histogram of ranks (middle column) and ECDF plots (right-hand side column) in the main text. (a) Model implementation is correct. (b) Parameter estimates are overdispersed relative to their true values. (c) Parameter estimates are underdispersed relative to their true values. (d) Parameter estimates are consistently overestimated relative to their true values. In the left-hand side column, the prior and replicate-averaged posterior (also known as the data-averaged posterior) distributions over some parameter *θ* are shown in light blue and dark blue, respectively. In the middle graphs, light-blue bands represent the 95%-confidence interval about the expected rank count, and horizontal black lines mark the rank count mean. Light-blue ellipses in the rightmost graphs represent confidence intervals about the empirical cumulative distribution function (ECDF).

Adherence to this distribution can be investigated by constructing histograms (Talts et al., 2018) as well as by looking at the empirical cumulative distribution function (ECDF) and their confidence bands (Säilynoja et al., 2021). When a model implementation fails RUV, it can do so in different ways. For instance, when the inference machinery leads to consistent overdispersed estimates, it produces a pattern of ranks concentrating around the middle rank (Fig. 6b). When underdispersion is present, on the other hand, ranks tend to bunch up towards the ends (Fig. 6c), creating a pattern of “horns”, which can also be caused by high autocorrelation in the MCMC draws. This is also why we recommend thinning MCMC draws in order to reduce autocorrelation. Figure 6d shows the rank patterns when the inference machinery produces biased estimates: ranks will bunch up against one of the ends, depending on whether estimates are biased downwards of upwards. In the particular case shown in figure 6c, the parameter at hand is being overestimated.

We conducted RUV on the three scenarios described in the previous section, which make use of the model depicted in Figure 1. In the interest of brevity, we only show the histograms and ECDFs for *R*, and leave the remaining plots for Λto the supplement (but see Box 1 below). As expected, under scenario 1 our model implementation passes the RUV – as indicated by histogram bars and ECDF values falling within their 95%-confidence intervals (Fig. 7a).

**Figure 7:**
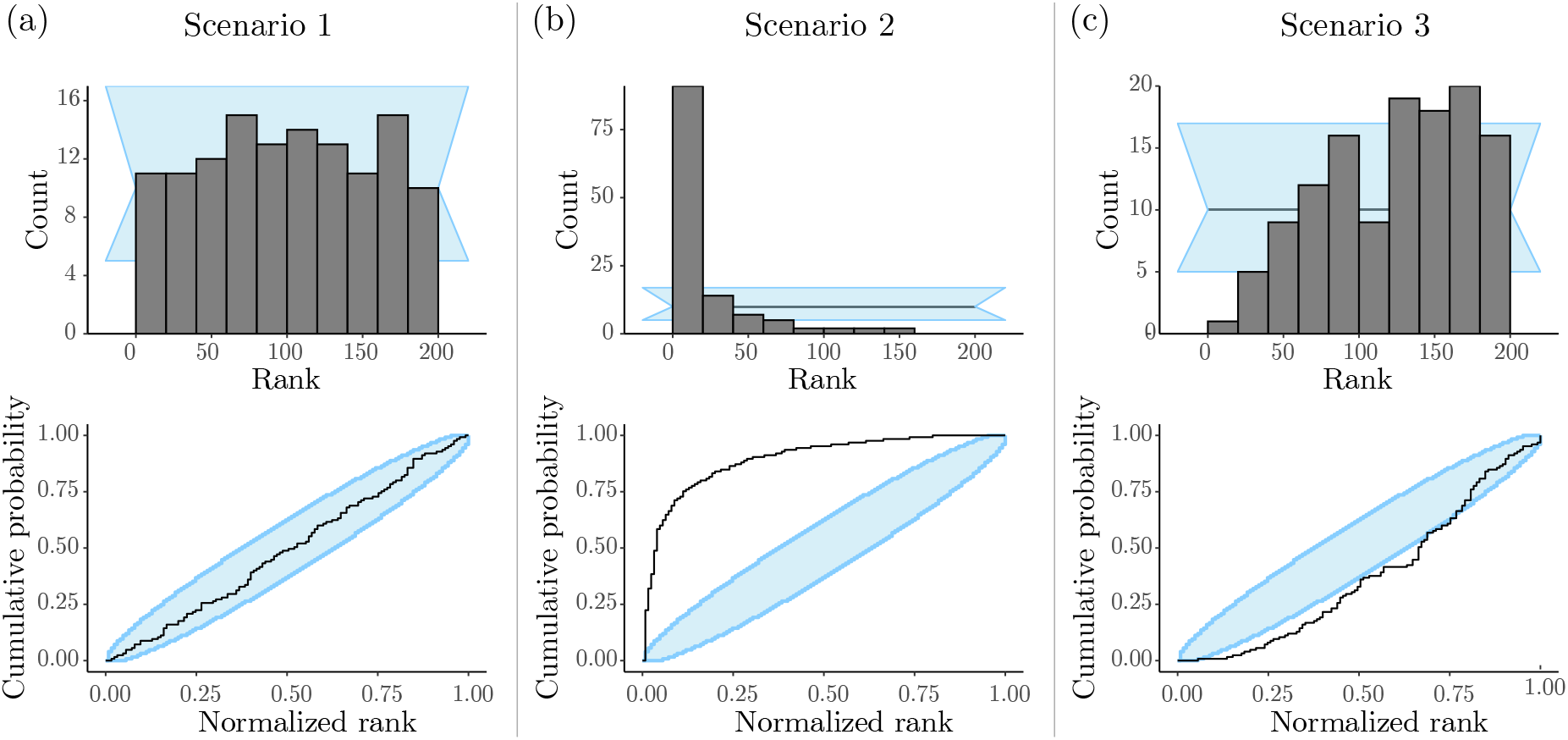
Rank-uniformity validation (RUV) of the Bayesian hierarchical model in Fig. 1. Panels in the top row show the histograms of *n* = 100 ranks, for parameter *R* in each scenario, obtained after 10% burnin and thinning of posterior samples down to 200 out of 10,000. Panels in the bottom row show the corresponding ECDF plots, for parameter *R* in each scenario. (a) In “Scenario 1”, the model was correctly specified and we can see that the ranks are compatible with a uniform distribution (within the blue band). (b) In “Scenario 2”, the inference machinery was misspecified and a clear pattern of overestimation shows up in the ranks, meaning the ranks for the data-generating values are usually smaller than expected under correctness. (c) In “Scenario 3” we can see a pattern of underestimation, evidenced by ranks bunching up to the right.

Under scenario 2, again as expected, our method failed RUV (Fig. 7b). In particular, we observes great overestimation of the Brownian motion rate (*R*). In a real world-analysis, these results would point to one or more faulty implementations (e.g., one or more model components, MCMC machinery, the simulator, etc). We remind the reader that in our experiment, scenario 2 was purposefully set up so that the (prior) models used in simulation and inference differed; our implementations are actually correct, but were induced to fail the RUV procedure.

Lastly, RUV results for scenario 3 contrasted with what we observed for this scenario’s coverage validation (Fig. 5c). While the model specified in scenario 3 passed its coverage validation (coverage was acceptable for both Λand *R*), it did not pass the RUV procedure. The corresponding rank histogram and ECDF plots indicate that *R* is underestimated (Fig. 7c). This result suggests that rank-uniformity validation can be more sensitive than coverage validation, at least for certain types of model misspecification, such as those affecting parameter estimate location.

Unlike scenario 2, in which we caused an explicit mismatch between the distributions used in simulation and inference, model misspecification under scenario 3 was subtler: simulation and inference models were identical (as in scenario 1), but tree samples from *f*_Φ_(*·*) (the Yule prior) were often rejected as ***θ*** was generated. The model used in scenario 3 failed RUV because rejecting tree samples induced an implicit Yule model in simulation that differed from the Yule model used in inference. Indeed, using a much simpler model and an analogous rejection scheme (Supplementary Fig. 2), we were able to recapitulate the results in Figure 7. While here we focus on so-called continuous parameters like *R* and Λ, it is also possible to conduct RUV to assess correctness on the space of phylogenies, a topic we leave for future research.

## Tree models

Tree models are stochastic processes that can capture the most fundamental tenet in evolutionary biology, namely common descent, at multiple time scales. Over the last few decades, pivotal theoretical work has not only characterized many properties of the more elementary tree models (for examples and an overview, see Nee, 2006; Wakeley, 2009; Stadler, 2013; Harmon, 2018), but also generalized them to be more realistic. Popular among empiricists, for example, are tree models that allow for lineage-affecting event rates that vary over time and across taxa, and that are state-dependent (“state” here meaning the attributes of a lineage’s genotypic, phenotypic, ecological or biogeographic characters). Such models lend themselves to the study evolutionary phenomena such as species diversification and infectious disease spread.

Although convenient evolutionary abstractions, tree models can nonetheless be challenging to formalize depending on their level of realism. The parameter space of a tree model is difficult to handle: it includes both a combinatorily complicated discrete component (the tree topology) and a continuous component (the branch lengths) (Semple et al., 2003). The theoretical properties, summarization, and exploration of tree space are all active topics of research in mathematical and computational biology (Gavryushkina et al., 2013; Gavryushkin and Drummond, 2016; Brown et al., 2020).

Given the interest in tree models shown by empirical, computational and theoretical biologists, in this section we will cover how tree models have been and can be validated, with an emphasis on tree space. We also propose a new manner by which tree models can undergo RUV and be assessed with respect to coverage. Our treatment is not meant to be an exhaustive review, but a short synthesis, and in keeping with the subject of the present work, we will not discuss protocols for the development and validation of Bayesian proposals in tree space. This topical subject is multifaceted (e.g., Douglas et al., 2021; Bouckaert, 2022; Douglas et al., 2022) and deserves a dedicated contribution we leave for the future.

One way a tree model implementation can be validated is by comparing statistical summaries of its samples (drawn through direct simulation or MCMC without data) against theoretical “target” values (e.g., Fig. 3). This type of validation can be compelling and is often easy to carry out, but closed-form expressions tend to be only available for simpler models like the birth-death process, the Kingman’s coalescent, and a few of their special cases and generalizations. Such expressions remain useful, nonetheless, as long as more complex models can be constrained to forms for which the relevant theory exists. Typical theoretical targets include the first moments of distributions on tree characteristics such as internal node ages, internal and terminal branch lengths, the sum of branch lengths, the number of tips, and the frequency of different tree topologies (Tajima, 1983; Rosenberg, 2002; Aldous and Popovic, 2005; Nee, 2006; Gernhard, 2006, 2008; Wakeley, 2009; Mooers et al., 2012).

As tree models increase in complexity (e.g., Maddison et al., 2007; Goldberg and Igić, 2012; Fitzjohn, 2010; Sciré et al., 2022), so does the tree space they define and as a consequence theoretical model validation as described above becomes difficult. When nodes can be serially sampled or direct ancestors of other nodes, for example, even enumerating all the possible trees under a model is a non-trivial exercise (Gavryushkina et al., 2013). The number of terminals in a tree can also complicate the theoretical characterization of tree models; except for when the number of terminals is small (Drummond and Bouckaert, 2015), algorithms have to be employed to generate expectations from theoretical principles (Kim et al., 2019).

When theoretical predictions useful for validation cannot be made, an often-employed method involves the comparison of independent model implementations, with one or both being simulators or inference engines. Tree samples from a direct simulator can be compared to samples drawn with MCMC without data (e.g., Zhang et al., 2023), or exact likelihood values can be compared between different implementations (Andréoletti et al., 2022). On extreme cases, however, even that strategy appears unattainable. For example, if it is unclear how to even directly simulate under a tree model – such as when node-age prior distributions are added to a birth-death process (models used in “node-dating”, Ho and Phillips, 2009; but see Heled and Drummond, 2012) – there seems to be no discernible path for validation in tree space.

The procedures of coverage validation and RUV can also be used to examine parameters in tree space. So far authors have mainly focused on the coverage of quantities such as species tree and gene root ages, sum of branch lengths, and number of direct ancestors (e.g., Gavryushkina et al., 2014; Ogilvie et al., 2022; Zhang et al., 2023). Fig. 8, for example, shows the coverage of the root age for the three scenarios we explored in the previous sections. Similarly to what was observed for tree-unrelated parameters, the root age had the expected coverage in scenarios 1 and 3, and RUV again aligned with the diagnosis of model misspecification for scenario 3.

**Figure 8:**
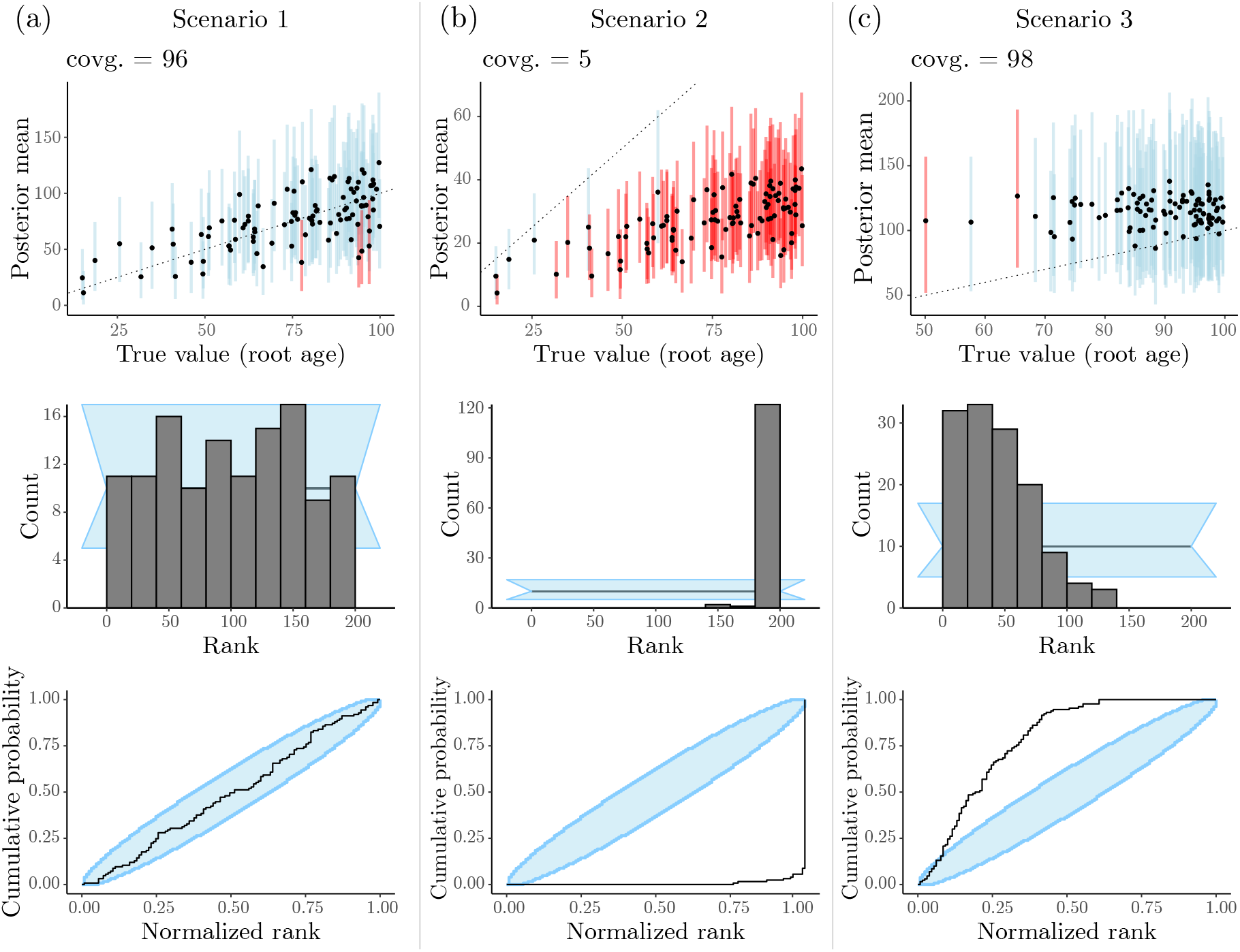
Coverage validation and rank-uniformity validation (RUV) of the Bayesian hierarchical model in Fig. 1, for each scenario described in the main text (also see Figs. 5 and 7), with respect to the height of *ϕ* (i.e., the phylogeny’s root age). Panels in the top row show the true (i.e., simulated) root age values plotted against their mean posteriors (the dashed line shows *x* = *y*). Dots and lines (100 per panel) represent true values and their 95%-HPDs, respectively. Simulations for which 95%-HPDs contained the true value are highlighted in blue, otherwise are presented in red. Panels in the middle row show the RUV histograms of *n* = 100 ranks in each scenario, obtained after 10% burnin and thinning of posterior samples down to 200 out of 10,000. Panels in the bottom row show the corresponding RUV ECDF plots in each scenario.

To the best of our knowledge, the RUV procedure has not been applied to validate phylogenetic models vis-a-vis tree space. This is likely because, due to the complexity of tree space, no canonical total-ordering structure (a requirement for ranking) has been proposed for trees. In what follows we fill this gap by introducing one possible solution for ranking trees as a pre-requisite for RUV.

Briefly, we propose that each of the trees in a set of MCMC samples, as well as the corresponding “true” tree, be first compared to an external, “reference” tree (see Algorithm 1 in the supplementary material). For example, a tree sample can be compared to the reference tree with respect to the length of their longest branch, to their topology, to their asymmetry, etc.; what matters here is that this comparison quantitatively measures the distance between the reference and the other tree. Then, once one knows how distant each true tree and its posterior samples are from the reference tree, they can be ranked relative to one another based on their associated distance measure. RUV proceeds normally from this point (see Box 2 for an example using the Robinson-Foulds distance; section 3 in the supplementary material illustrates RUV and coverage validation with other distance metrics).

### Box 2

**Validating a phylogenetic model with respect to its phylogenetic tree parameter**

Given the centrality of the phylogenetic tree (Φ) in comparative analyses, one must pay close attention to this parameter when validating a phylogenetic model. This is no easy task: tree space is defined by a complex mix of a discrete and continuous components, and one in which trees cannot be easily ranked (see main text).

To get around this difficulty, we propose computing one or more phylogenetic metrics, or functionals, for which total-ordering holds and for which ranks can thus be obtained. One such metric is the well-known Robinson-Foulds distance between two trees (RF; Robinson and Foulds, 1981), which counts the number of clades implied by one tree but not the other. In order to compute a relational metric like the RF distance during validation, we must have a reference phylogeny *ϕ*_0_ to which we can compare our focal generating phylogeny *ϕ* and its posterior MCMC samples. The RUV protocol remains the same, with an additional step in which we generate *ϕ*_0_ (see Algorithm 1 in the supplementary material). Fig. 9 shows validation results for the RF metric for five-taxon trees simulated under a simple Kingman’s coalescent model assuming a known effective population size of 1.0.

**Figure 9:**
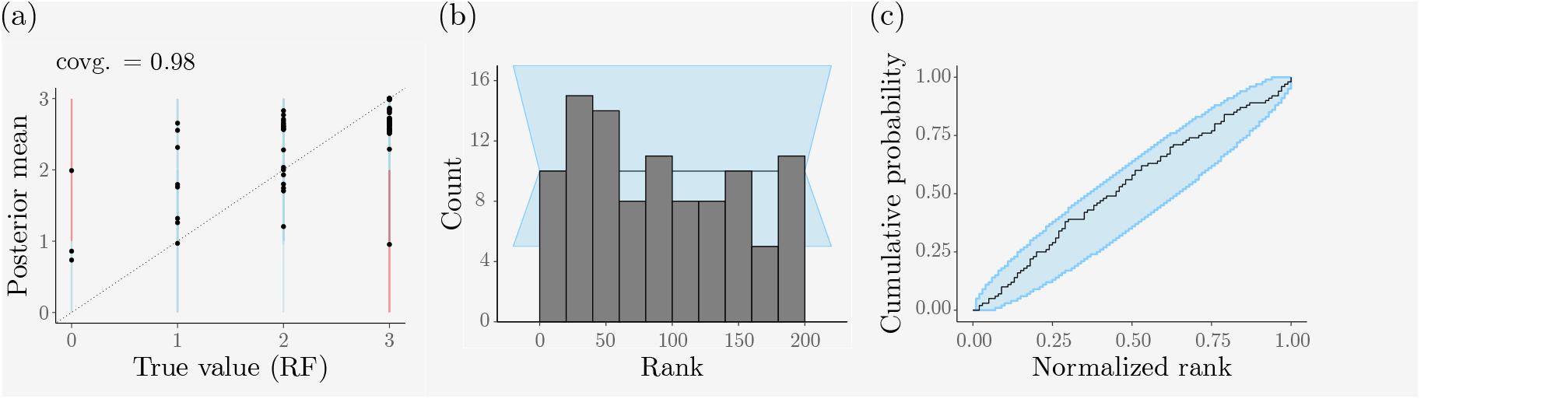
Coverage validation and rank-uniformity validation (RUV) of a Kingman’s coalescent model with respect to the Robinson-Foulds distance (RF; see text in box) between the coalescent tree (Φ) and a reference (random) tree (*ϕ*_0_). The effective population size parameter is assumed known and fixed to 1.0 during inference. (a) The true RF distances (i.e., between 100 simulated coalescent trees, ***ϕ*** =*{ϕ*_*i*_ : 1 *≤ i ≤* 100*}*, and the same reference tree, *ϕ*_0_) plotted against the corresponding mean posterior RF distances (calculated from posterior samples of each *ϕ*_*i*_ and *ϕ*_0_). The dashed line shows *x* = *y*. Dots and vertical lines represent true RF-distance values and their estimated 95%-HPDs, respectively. Simulations for which 95%-HPDs contained the true value are highlighted in blue, otherwise are presented in red. (b) RUV histograms of 100 ranks obtained after 10% burnin and thinning of posterior samples down to 1000 out of 10, 000. (c) RUV empirical CDF plots.

Fig. 9a shows the coverage of 95%-HPD intervals of the RF distance metric, while Fig. 9b-c give its rank distribution and empirical CDF, respectively. The coverage of the RF metric is very close to 95 (see Table 3, with *n* = 100), and the rank distribution is approximately uniform on (1, *L* + 1); together, these panels indicate this model is correctly implemented. Note that for large numbers of species, *ϕ*_0_ is unlikely to share internal nodes with *ϕ*’s posterior samples (Steel, 1988), in which case the RF distance metric may not be so useful. In the supplement we consider other phylogenetic tree metrics that could be used as an alternative or in addition to the RF distance (Supplementary Fig. 4).

## Software

We implemented a suite of methods for automating many of the steps involved in coverage validation and RUV. These methods were developed in Java and integrated into the BEAST 2 platform (Bouckaert et al., 2019). Code and a tutorial are available on https://github.com/rbouckaert/DeveloperManual.

## Bayesian model validation guidelines for developers and reviewers

In the previous sections, we described and executed two procedures for validating Bayesian models, namely, coverage validation and rank-uniformity validation (RUV). Once both simulation and inference can be conducted under a model *ℳ*, executing these procedures amounts to following relatively straightforward protocols (Fig. 4). Importantly, following these protocols should validate any Bayesian model, regardless of the nature of its parameter space and its component sampling distributions. This is because such protocols provide clear, objective rules for assessing model correctness, based on the coverage and rank distribution of parameters values; both can be computed for any and all Bayesian models. For the reasons above, carrying out an analysis of coverage and/or of the distribution of parameter-value ranks (with respect to their posterior samples) should on one hand be a requirement, and on another should suffice for introducing a new Bayesian model implementation.

### My model implementation failed correctness tests, what now?

Method developers should expect their software to often fail validation, especially at early development stages, causing the loops in Fig. 4 to be visited many times. The validation procedure is almost always arduous and repetitive, but very effective in revealing issues and in giving modelers peace of mind when releasing their software. A correctly implemented inference machinery can nonetheless still fail a validation test if there is some unforeseen form of model misspecification (e.g., truncation, see scenarios in Figure 5). In such cases, a potentially delicate stage of method development begins, when decisions must be made between further testing or software release.

If validation success is marginal or contingent upon a substantially constrained parameter space, or if a Bayesian method has good coverage but fails the demanding RUV (as shown here and elsewhere; McHugh et al., 2022), further simulation experiments might illuminate the nature of the model misspecification and suggest ways to modify the model. For example, developers may want to tweak an aspect of simulation, and then repeat RUV in search for regularities in parameter over- or underestimation (e.g., section 3 in the supplement). When releasing a method despite validation failure, researchers should in the very least be expected to report all attempts made to validate an implementation, why they seemed to fail, and what biases were uncovered, if any. Ideally, guidelines should be provided for interpreting results obtained with a tool known to be biased.

When confronted with utter validation failure, we urge method developers to resist the temptation of downplaying the importance of the validation effort, and instead ask the hard question of whether their models are reasonable in the first place. If large numbers of simulations must be rejected so as to obtain realistic data – or data whose probabilities can be calculated – this could be a sign that the model needs to be modified. Independent implementations that also do not pass validation tests provide further evidence that the issue is potentially in the model assumptions themselves. Historically, model design has often gone in the direction of incrementally conditioning the statistical process so as to match empirical observations (e.g., Gelman and Meng, 1996; Gelman et al., 2020), but re-imagining the model entirely might be the best solution.

### Model characterization

In addition to the model validation we detailed above, there is an infinite number of ways in which a new or published model can have its behavior inspected. Researchers may want to know, given a model, how sensitive parameter estimates are to data set size, prior choice, model complexity, violation of model assumptions, to name a few. Studies have examined how these factors affect estimation accuracy and precision (e.g., Zhang et al., 2023; Luo et al., 2023), as well as the mixing and convergence of MCMC chains (e.g., Nylander et al., 2004; Zhang et al., 2023). We collectively refer to these examinations as “model characterization”: any analysis of model behavior beyond assessing its correctness. Model characterization is rarely carried out to satisfy the curiosity of the theoretician (but see, e.g., Tuffiey and Steel, 1997; Steel and Penny, 2000); it is instead normally motivated by a model’s empirical applications. These investigations are thus critical for the longevity and popularity of a model, as domain experts will only adopt a model widely if they know when to trust the results and how to interpret them.

It is possible to characterize certain aspects of model behavior while simultaneously verifying its correctness, as discussed in the coverage validation section above. For example, one can observe how accurate parameter estimates are (e.g., if the points in Fig. 5 fall on the identity line) under both correctly and incorrectly specified models. However, the requirement of simulating parameter values from a prior distribution *f*_Θ_(*·*) during the validation of a model can complicate its characterization. Depending on the characterization experiment’s goals and design, researchers may find themselves rejecting a large fraction of simulated data sets – perhaps because data sets do not resemble those in real life, or because they are too large to analyze. But as we showed, rejecting draws in simulation may then be picked up by the validation protocol as an incorrectly implemented model. This problem can only worsen the more dimensions of parameter space are allowed to vary. In most cases, it may thus make more sense to first verify model correctness by following the procedures we described above, and then characterize model behavior further in a subsequent batch of analyses.

We conclude this section by proposing that scientists contributing or reviewing a new model ask the following question: Is the contribution at hand carrying out an empirical analysis that will specifically profit from scrutinizing model behavior? If not, then model characterization efforts will likely serve their purpose better elsewhere, and profit from being shouldered by the scientific community at large.

### Concluding remarks

In order to keep up with the large amounts of data of different kinds accumulating in public databases, researchers in the life sciences must constantly update their computational tool boxes. New models are implemented in computational methods every day, but if they are not properly validated, downstream conclusions from using those methods may be void of any significance.

In the present study, we described and executed two distinct validation protocols that verify a Bayesian model has been correctly implemented. Although we looked at examples from evolutionary biology, specifically statistical phylogenetics, these two simulation-based protocols work for any and all Bayesian models.

We further elaborate on the difference between experiments in model validation versus model characterization. Newly implemented models can only profit from validation experiments, which are strictly concerned with theoretical expectations (e.g., about coverage) a model must meet if correctly implemented. Model characterization, on the other hand, is about inspecting model behavior as a variety of data set and model attributes interact; here, exact quantitative predictions may not be theoretically guaranteed. Such experiments are best designed and justified when empirically motivated.

We hope the guidelines described here can enhance both the release rate and standards of statistical software for biology, by assisting its users, developers and referees in quickly finding common ground when evaluating new modeling work.

