## Supplementary material for "How to validate a Bayesian evolutionary model"

February 7, 2024

### Supplementary Material

### 1 Validating a phylogenetic Brownian motion simulator

In this section we focus on validating a simulator for the phylogenetic Brownian motion model ("PhyloBM"; **?**). As explained in the main text, our goal is to verify that the expected value of certain summary statistics (given a specific combination of parameter values) falls within its $\alpha$-confidence intervals approximately $\alpha$% of the time. We will build confidence intervals about statistics calculated from several PhyloBM samples of size 10,000, and then ask if the "population" value of a statistic – given by the parameters of the multivariate normal sampling distribution – is contained within its confidence interval frequently enough.

For summary statistics, we pay attention to the trait value's (i) species mean, (ii) species variance, (iii) among-species covariance, and (iv) among-species correlation coefficient. Supplementary figure 1 shows one hundred confidence intervals for each of these statistics, under multivariate normal $\text{MVN}(\boldsymbol{y_0}, r\boldsymbol{T})$, where $\boldsymbol{y_0} = \{0, 0, 0\}$, $r = 0.1$ and $\boldsymbol{T}$ is given by the tree in Fig. 2 in the main text. Supplementary table 1 summarizes how often each statistic fell within its 95%-confidence interval. These results indicate the PhyloBM simulator produces appropriate confidence intervals and behaves as expected.

**Supplementary Table 1:** The number of times $k$ that a summary statistic was contained within its corresponding 95%-confidence interval. Each statistic was calculated from 100 datasets of size 10,000 simulated under the PhyloBM model described in the text and in Box 1 in the main text.

| Statistic | Species $s$ (and $v$) | $k$ |
|---|---|---|
| $\text{E}[y_s]$ | A | 95 |
| | B | 97 |
| | C | 98 |
| $\text{Var}[y_s]$ | A | 93 |
| | B | 97 |
| | C | 97 |
| $\text{Cov}[y_s, y_v]$ | A and B | 95 |
| | A and C | 95 |
| | B and C | 97 |
| $\text{Cor}[y_s, y_v]$ | A and B | 96 |
| | A and C | 96 |
| | B and C | 97 |

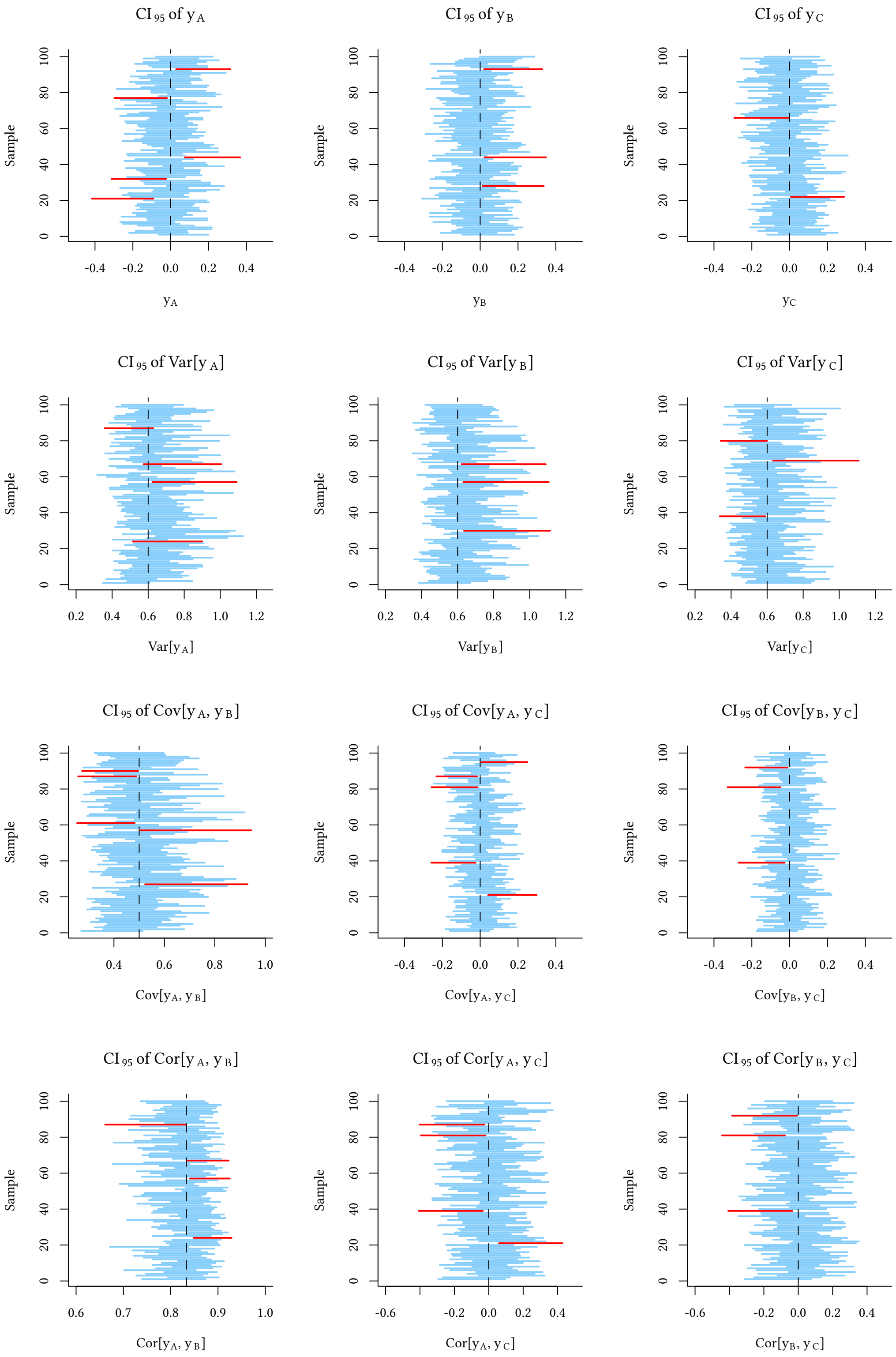


**Supplementary Figure 1:** One hundred 95%-confidence intervals (blue and red lines) built for four different summary statistics, when validating a phylogenetic Brownian motion model simulator. Red lines represent intervals that do not contain the value expected under the MVN sampling distribution defined by a bifurcating three-taxon phylogenetic tree. Summary statistics include each leaf's character-value mean (top row) and variance (second row from the top), as well as pairwise (leaf) character-value co-variances (third row from the top) and correlations (bottom row).

#### 2 Model validation with rejection sampling: a simple example

In this section, we experiment with a simple hierarchical Gaussian (toy) model to further examine the effect of rejection sampling in coverage validation and rank-uniformity validation (RUV). This experiment is motivated by results described in the main text, namely, the model in scenario 3 (Fig. 1, with extreme rejection sampling) passing coverage validation but not RUV.

Let us devise the following data-generating process for obtaining different levels of model misspecification:

$$\mu \sim \text{Normal}\,(0,1)\; T[t,],$$
$$y_1,\ldots,y_K \overset{\text{iid}}{\sim} \text{Normal}\,(\mu,1)\,,$$

where $K$ is the sample size, and notation $T[,t]$ indicates the distribution is truncated **below** at $t \in \mathbb{R}$, i.e.,

$$\pi_t(\mu) = \frac{\phi(\mu)}{1-\Phi(t)}\mathbb{I}(\mu > t).$$

and $\phi$ and $\Phi$ are the pdf and CDF of a standard normal, respectively. Here, $\mu$ and $y_1,\ldots,y_K$ correspond to $\theta_i$ and $d_i$ in Fig. 4 (main text), respectively.

Inference is then done under the misspecified model

$$\mu \sim \text{Normal}\,(0,1)\,,$$
$$y_1,\ldots,y_K \overset{\text{i.i.d.}}{\sim} \text{Normal}\,(\mu,1)\,.$$

It is well-known that:

$$\mu \mid y_1,\ldots,y_K \sim \text{Normal}\left(\frac{\sum_{k=1}^{K} y_k}{K+1}, \frac{1}{K+1}\right), \tag{1}$$

i.e., the posterior is known in closed-form so MCMC is not required and one can sample directly and independently from it. One can thus control how extreme the model misspecification will be during inference by increasing $t$. Here we explore $t = \{0, 1, 1.5\}$ and $K = \{5, 10, 50\}$ in order to show how the method behaves in various misspecification scenarios. We ran $n = 1000$ replications of the experiment, draw 200 i.i.d. samples directly from the posterior in (1).

The resulting rank histograms (Supplementary Fig. 2) show how the posterior consistently underestimates the true mean, as indicated by a pattern of ranks bunching up towards the right-hand side of the histogram. The pattern is clearer in situations where the evidence in the data is weaker (e.g., when $K$ is small), and the effect of the prior on the posterior is stronger. Moreover, the misspecification becomes more apparent the larger $t$ is, that is, the more extreme the misspecification becomes.

It is worth noting a couple of things. First, unlike what we observed and reported for the model in the main text, coverage validation does suggest an incorrectly specified model for certain combinations of $t$ and $K$ (Supplementary Table 2). Second, when the truncation is less extreme ($t = 0$) and there is a substantial amount of data ($K = 50$), RUV fails to detect any problems. This is an important insight about validation protocols: their sensitivity is context-dependent, and responds to the combination of model and data regime.

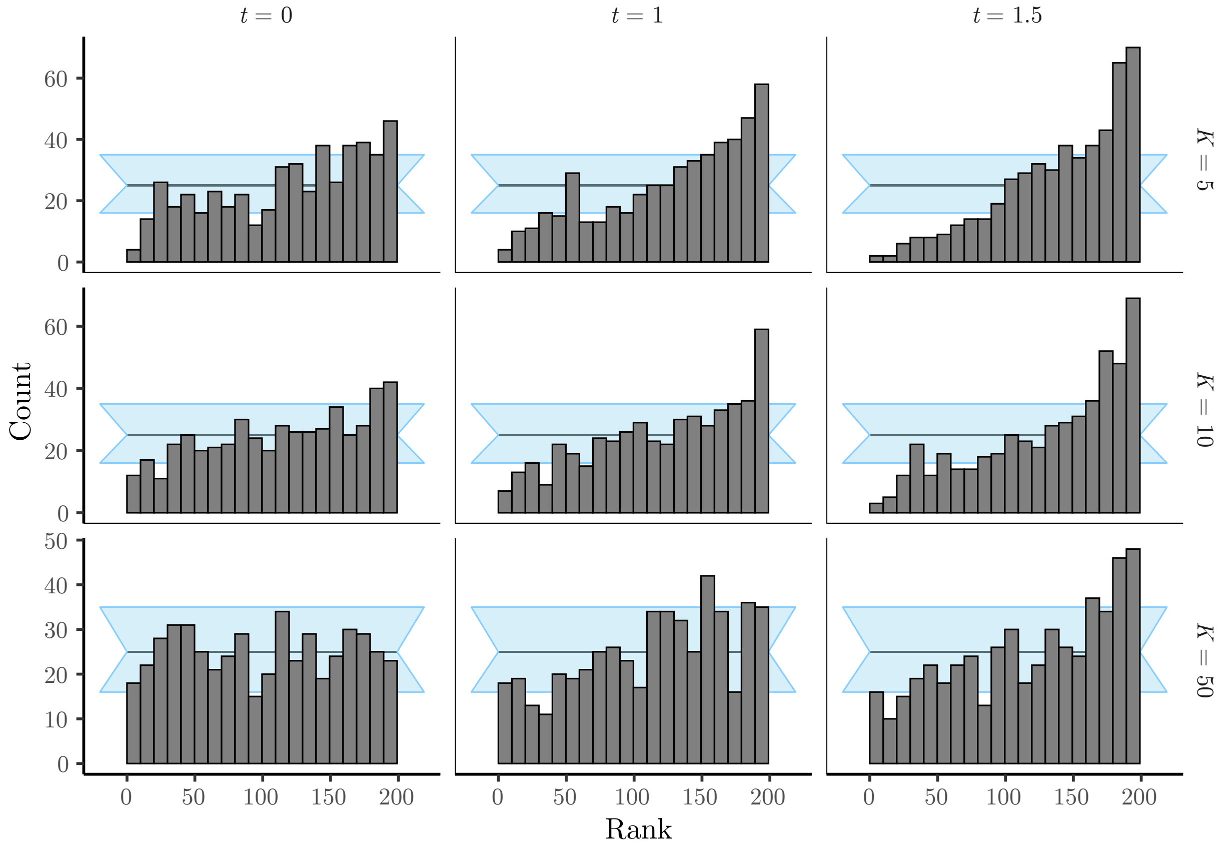


**Supplementary Figure 2:** Rank-uniformity validation (RUV) of the hierarchical Gaussian toy model. We show the rank histograms in three misspecification scenarios ($t$ is 0.0, 1.0 or 1.5) and three data regimes ($K$ is 5, 10 or 50). Results are based on $n = 1000$ replicates of $L = 200$ i.i.d. draws each. Horizontal black lines shows the expected count for each rank (the same for all ranks due to uniformity), and the light-blue bands represent the 95%-confidence interval for the counts.

**Supplementary Table 2:** Coverage results for the 95%-HPD under the hierarchical Gaussian toy model. For each combination of truncation ($t$) and data set size ($K$), we show the estimated coverage over $n = 1000$ replicates, and whether the coverage procedure passes. A pass is determined according to table 2 in the main text.

| Truncation ($t$) | Number of observations ($K$) | Estimated coverage | Pass |
|---|---|---|---|
| 0 | 5 | 0.93 | No |
| 1 | 5 | 0.92 | No |
| 1.5 | 5 | 0.90 | No |
| 0 | 10 | 0.95 | Yes |
| 1 | 10 | 0.91 | No |
| 1.5 | 10 | 0.91 | No |
| 0 | 50 | 0.96 | Yes |
| 1 | 50 | 0.93 | No |
| 1.5 | 50 | 0.93 | No |

### 3 Validating a phylogenetic model with respect to a phylogenetic tree parameter, $\Phi$

Our goal is to evaluate if an inferential implementation of model $\mathcal{M}$, $\mathrm{I}[\mathcal{M}]$, is well-calibrated and correct. In this section, we will focus on one critical parameter of $\mathcal{M}$: the phylogenetic tree $\Phi$. We will pay special attention to $\Phi$ while carrying out coverage validation and the RUV procedure. In practice, this amounts to computing the coverage of $\Phi$ and the rank distribution of this parameter in comparison to its posterior samples.

What feature of $\Phi$ should one use when calculating coverage and ranks? Because of the nature of tree space, summarizing and comparing phylogenetic trees with univariate measures is not a trivial task. The key to an effective validation effort is to choose a functional that reflects relevant estimators of the quantity of interest, in our case, $\Phi$. Fortunately, it is possible to exploit the metric nature of tree space and compute quantities both from a sampled phylogenetic tree $\phi$ – i.e., the "true" tree simulated during the validation process – as well as distances with respect to a reference phylogeny $\phi_0$. We summarize some of these distances in supplementary table 3.

**Supplementary Table 3:** Metrics (functionals) for investigating model correctness in phylogenetic tree space. $\phi$ is a (phylogenetic tree) sample of $\Phi$, sampled from a tree model (e.g., a Yule process). $\phi_0$ is a reference tree to which $\phi$ and its posterior samples (see Algorithm 1 and the RUV section in the main text) are being compared, with respect to any of the metrics in the table. For the KC metric, the parameter ($\omega$) controls the balance between topological and branch length information and was set at $\omega = 1/2$.

| Metric | Notation | Ref. |
|---|---|---|
| The largest branch length in $\phi$ | $\mathrm{LB}(\phi)$ | N/A |
| The length of $\phi$ (the sum of all branch lengths) | $\mathrm{LEN}(\phi)$ | N/A |
| The difference between the largest and smallest branch length of $\phi$ | $\mathrm{R}(\phi)$ | N/A |
| The Robinson-Foulds distance between $\phi$ and $\phi_0$ | $\mathrm{RF}_0(\phi)$ | (Robinson and Foulds, 1981) |
| The Kendall-Coljin distance between $\phi$ and $\phi_0$ | $\mathrm{KC}_0(\phi;\omega)$ | (Kendall and Colijn, 2016) |
| The Billera-Holmes-Vogtman distance between $\phi$ and $\phi_0$ | $\mathrm{BHV}_0(\phi)$ | (Billera et al., 2001) |

In the case of tree-space distance metrics, we must slightly modify our RUV procedure (Algorithm 1). The key difference is the sampling of a reference phylogenetic tree, $\phi_0$, prior to the simulation of the $n$ i.i.d. $\Phi$ samples,

$\boldsymbol{\phi} = \{\phi_i : 1 \leq i \leq n\}$. Given a reference tree $\phi_0$, each sampled $\phi_i$ and all of its $L$ posterior MCMC samples are then compared to $\phi_0$ with respect to a chosen distance metric. It is the evaluated distance metric that will underlie the ranking of $\phi_i$ relative to its posterior MCMC samples. Once ranks are computed, RUV proceeds as usual.

**Algorithm 1:** Algorithm for carrying out a rank-uniformity validation procedure with respect to the phylogenetic tree parameter $\Phi$. Parameters $\boldsymbol{\theta}$ include both the tree parameter, $\theta_\Phi$, and non-tree (scalar) parameters, $\boldsymbol{\theta}_s$. Data $\boldsymbol{d}$ represents the output of an evolutionary process taking place along the phylogenetic tree (e.g., a continuous-time Markov chain modeling DNA substitutions).

```
n ← 100
                                         /* Number of data sets to simulate */
θ_H ← InitializeHyperparameters()
                             /* Hyperparameters initialized to constant values */
θ_0 ← SampleNonTreeParameters(θ_H)
                                   /* θ_0 ~ f_Θ(·), with θ_0 = {θ_Φ,0, θ_s,0} */
φ_0 ← SampleTree(θ_Φ,0)
                                           /* φ_0 ~ f_Φ|Θ_Φ(·|Θ_Φ = θ_Φ,0) */
for i ← 1 to n do
    θ_i ← SampleNonTreeParameters(θ_H)
                                   /* θ_i ~ f_Θ(·), with θ_i = {θ_Φ,i, θ_s,i} */
    φ_i ← SampleTree(θ_Φ,i)
                                           /* φ_i ~ f_Φ|Θ_Φ(·|Θ_Φ = θ_Φ,i) */
    d_i ← SampleData(φ_i, θ_s,i)
                                /* d_i ~ f_D|Φ,Θ_s(·|Φ = φ_i, Θ_s = θ_s,i) */
    θ'_i ← MCMC(f_D|Θ(d_i|Θ = θ_i) f_Θ(θ_i))
          /* θ'_i = {θ_i^j : 1 ≤ j ≤ L}, where L is the number of MCMC samples */
    δ̄_i ← CalculateDistance(φ_0, φ_i)
                                  /* According to a distance metric of choice */
    δ_i^j ← CalculateDistance(φ_0, φ_i^j)
                                             /* δ_i = {δ_i^j : 1 ≤ j ≤ L} */
    r_i ← GetRank(δ̄_i, δ_i^j)
                                       /* r_i = Σ_{j=1}^{L} 𝕀(δ_i^j < δ̄_i) */
end
if IsRankDistributionUniform(r) then
    return true
end
else
    return false
end
```

Supplementary figures 3 and 4 summarize the coverage and RUV results, respectively, for the metrics listed in supplementary table 3, when the model is the simple Kingman's coalescent. The model consists of a single five-taxon phylogenetic tree parameter, $\Phi$, assuming a known effective population size of 1.0.

We can first verify that our model implementation is well-calibrated with respect to the phylogenetic tree parameter, as shown by appropriate 95%-HPD-coverage statistics over the different tree metrics (Supplementary Fig. 3). The evidence for implementation correctness is enhanced by observing that the ranks for the different metrics

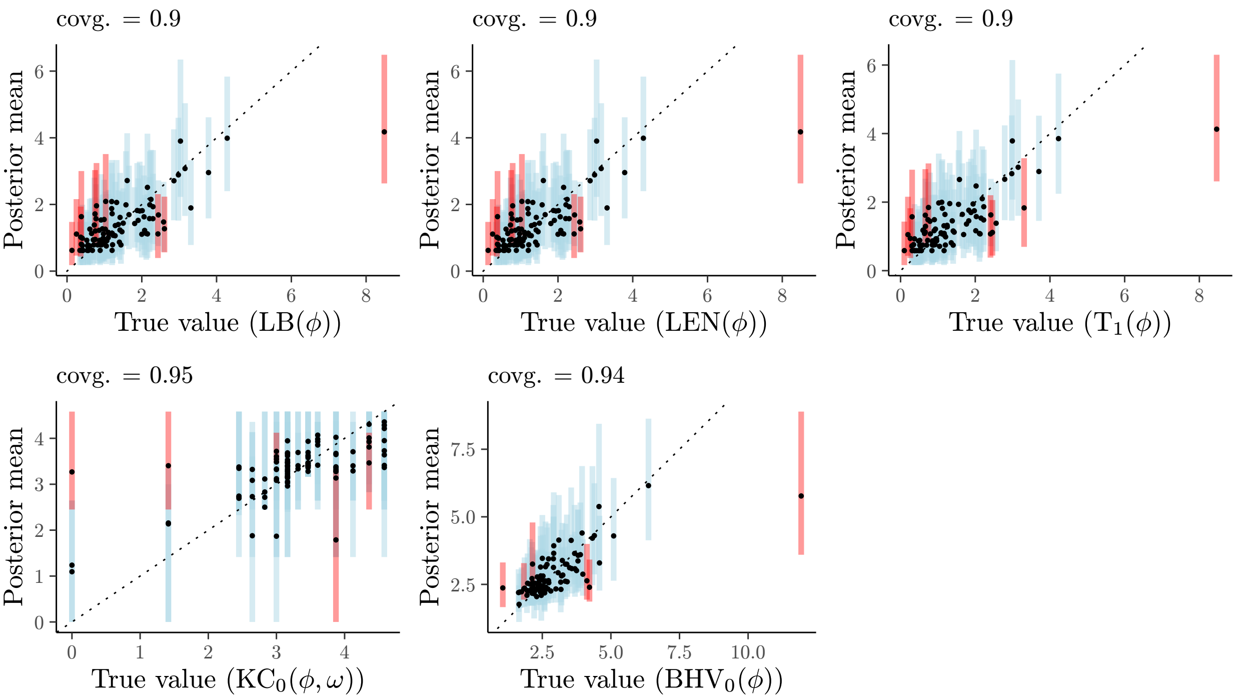


**Supplementary Figure 3:** Coverage validation of a phylogenetic model, focusing on the phylogenetic tree parameter (see Supplementary Table 3 for a description of the different distance metrics).

are (approximately) uniformly distributed, lying inside their confidence bands (Supplementary Fig. 4a), and their corresponding ECDFs also lie well inside their confidence ellipses (Supplementary Fig. 4b).

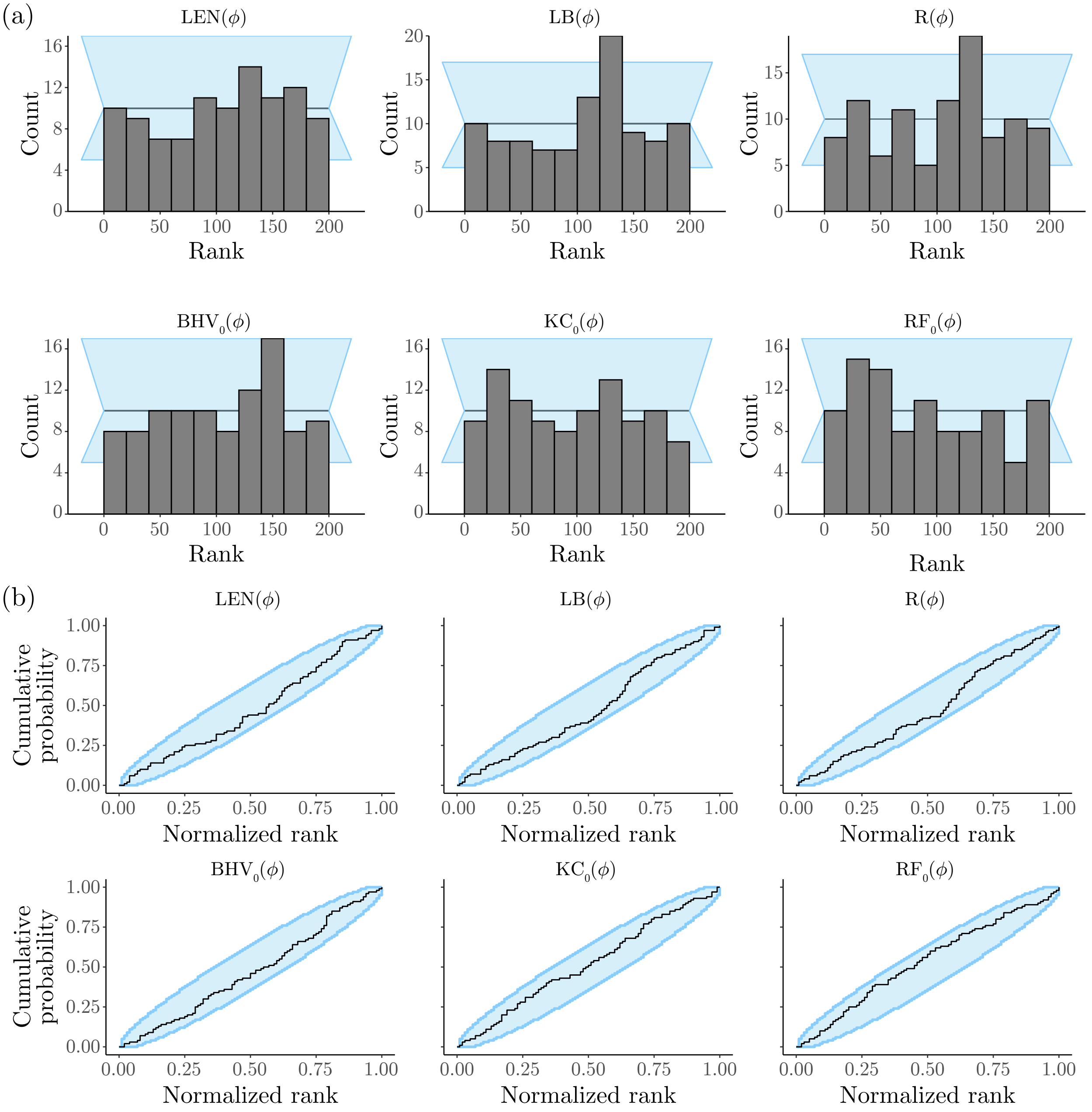


**Supplementary Figure 4:** Rank-uniformity validation (RUV) of a phylogenetic model, focusing on the phylogenetic tree parameter (see Supplementary Table 3 for a description of the different distance metrics). (a) Rank distribution for each metric. (b) Empirical cumulative distribution function (ECDF) for each metric.

### 4 Proof for coverage validation

In this section we provide a mathematical argument that coverage-based validation is sound, i.e., that sampling from the prior, simulating data and then using the same prior for computing the posterior should give Bayesian credible intervals (BCIs) with nominal frequentist coverage.

For a number $n$ of replicates, simulate parameter values $\theta_i$, and then given those values, simulate data $d_i$:

$$\theta_i \sim f_{\Theta}(\cdot),$$
$$d_i \sim f_{D|\Theta}(\cdot|\Theta = \theta_i).$$

Now for notational convenience, define $a_i := a(d_i, \alpha)$ as the HPD lower bound and, similarly, $b_i := b(d_i, \alpha)$ as the HPD upper bound. Recall $I_\alpha(d_i)$ is such that:

$$Q_{d_i}(b_i) - Q_{d_i}(a_i) = p_1 - p_2 = \alpha,$$

where $Q_d(x)$ is the posterior CDF (conditional on data $d$) and $p_1, p_2 \in (0, 1)$, with $p_1 < p_2$. A natural quantity to compute is:

$$S_n = n^{-1} \sum_{i=1}^{n} \mathbb{I}\left(\theta_i \in I_\alpha(d_i)\right),$$

i.e., the attained coverage of the Bayesian intervals.

Let $F_U(x) = x$ be the CDF of a $\text{Uniform}(0, 1)$ random variable. Now we can consider what happens when the number of simulations grows, i.e., the limit $\lim_{n\to\infty} S_n$. We may re-write the limit as:

$$\begin{aligned}
\lim_{n\to\infty} S_n &= \lim_{n\to\infty} n^{-1} \sum_{i=1}^{n} \mathbb{I}\left(\theta_i \in I_\alpha\left(d_i\right)\right), \\
&= \lim_{n\to\infty} n^{-1} \sum_{i=1}^{n} \left\{\mathbb{I}\left(\theta_i \leq b_i\right) - \mathbb{I}\left(\theta_i \leq a_i\right)\right\}, \\
&= \lim_{n\to\infty} n^{-1} \sum_{i=1}^{n} \mathbb{I}\left(\theta_i \leq b_i\right) - n^{-1} \sum_{i=1}^{n} \mathbb{I}\left(\theta_i \leq a_i\right), \\
&= \lim_{n\to\infty} n^{-1} \sum_{i=1}^{n} \mathbb{I}\left(Q_{d_i}^{-1}\left(\theta_i\right) \leq p_1\right) - \lim_{n\to\infty} n^{-1} \sum_{i=1}^{n} \mathbb{I}\left(Q_{d_i}^{-1}\left(\theta_i\right) \leq p_2\right), \\
&= F_U(p_1) - F_U(p_2) = \alpha,
\end{aligned}$$

where the last line follows from the fact that the CDF of $\theta_i$ is uniformly distributed on $(0,1)$ (Theorem 1 in Cook et al., 2006) and almost sure convergence of the ECDF to the true CDF due to the Glivenko-Cantelli theorem (Billingsley, 1986, page 275).

### 5 Other supplementary figures

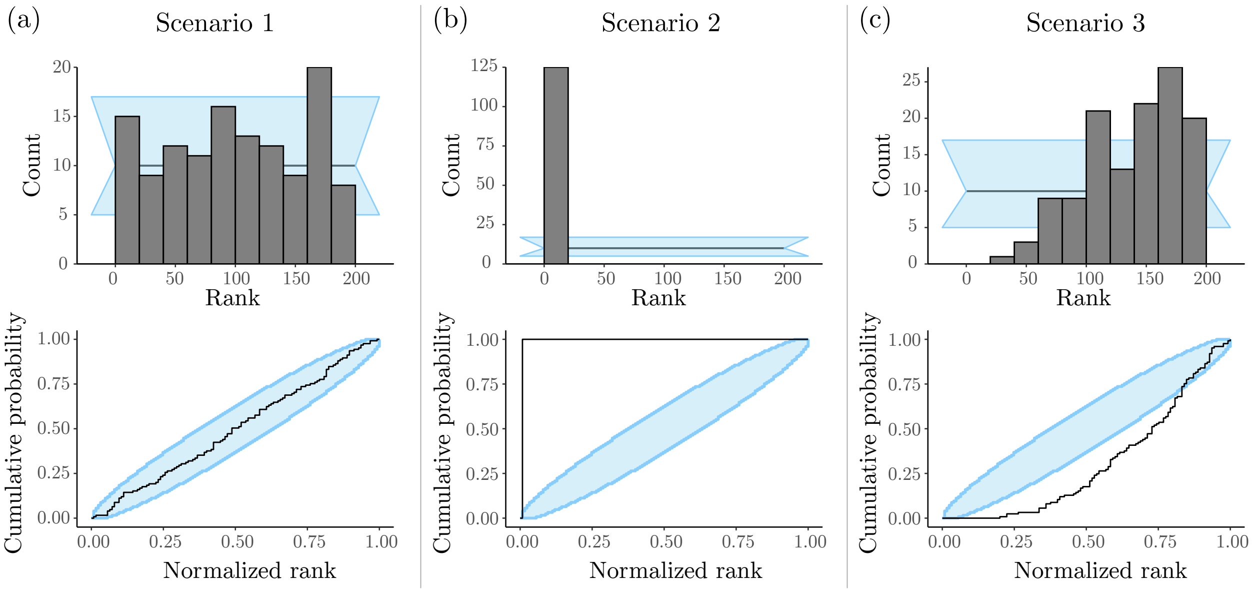


**Supplementary Figure 5:** Rank-uniformity validation (RUV) of the Bayesian hierarchical model in Fig. 1 in the main text. Panels in the top row show the histograms of $n = 100$ ranks, for parameter $\Lambda$ in each scenario, obtained after 10% burnin and thinning of posterior samples down to 200 out of 10,000. Panels in the bottom row show the corresponding ECDF plots, for parameter $\Lambda$ in each scenario. (a) In "Scenario 1", the model was correctly specified, and we simulated trees with 3 to 300 taxa using rejection sampling (approximately one in ten trees were rejected). (b) In "Scenario 2", the model was incorrectly specified in inference (see main text), and we used the same data sets simulated in "Scenario 1". (c) In "Scenario 3", the model was correctly specified, but rejection sampling was more intense (we rejected a large number of trees, approximately 90%, keeping those having between 100 to 200 tips).
